# Sensory coding and contrast invariance emerge from the control of plastic inhibition over emergent selectivity

**DOI:** 10.1101/2020.04.07.029157

**Authors:** René Larisch, Lorenz Gönner, Michael Teichmann, Fred H. Hamker

**Affiliations:** TU Chemnitz, Dept. of Computer Science, Artificial Intelligence; TU Dresden, Faculty of Psychology, Lifespan Developmental Neuroscience; Bernstein Center Computational Neuroscience Berlin

## Abstract

Visual stimuli are represented by a highly efficient code in the primary visual cortex, but the development of this code is still unclear. Two distinct factors control coding efficiency: Representational efficiency, which is determined by neuronal tuning diversity, and metabolic efficiency, which is influenced by neuronal gain. How these determinants of coding efficiency are shaped during development, supported by excitatory and inhibitory plasticity, is only partially understood. We investigate a fully plastic spiking network of the primary visual cortex, building on phenomenological plasticity rules. Our results suggest that inhibitory plasticity is key to the emergence of tuning diversity and accurate input encoding. We show that inhibitory feedback (random and specific) increases the metabolic efficiency by implementing a gain control mechanism. Interestingly, this led to the spontaneous emergence of contrast-invariant tuning curves. Our findings highlight that (1) interneuron plasticity is key to the development of tuning diversity and (2) that efficient sensory representations are an emergent property of the resulting network.

## 1 Introduction

The primary visual cortex (V1) represents visual stimuli in a highly efficient manner (Froudarakis et al., 2014; Dadarlat & Stryker, 2017). Recent research has identified two distinct factors underlying the efficiency of visual representations: First, representational efficiency in terms of absolute information content, which is mainly determined by the receptive field tuning diversity (Goris et al., 2015). Second, metabolic efficiency in terms of the number of spikes required to represent a specific input stimulus. This aspect is strongly influenced by gain control mechanisms caused by inhibitory feedback processing (Carvalho & Buonomano, 2009; Isaacson & Scanziani, 2011). How these determinants of coding functionality are shaped is only partially understood. While it has long been known that excitatory plasticity is necessary for the development of an accurate and efficient input representation (Olshausen & Field, 1996; Bell & Sejnowski, 1997; Zylberberg et al., 2011), there has recently been growing interest in the role of inhibitory plasticity, fueled by recent studies demonstrating plasticity at inhibitory synapses (Khan et al., 2018). As the synaptic plasticity of inhibitory interneurons in V1 likely exerts strong effects on the outcome of excitatory plasticity (Wang & Maffei, 2014), complex circuit-level interactions occur between both types of plasticity. This notion has received further support based on recent theoretical studies (Mongillo & Loewenstein, 2018). Above all, these findings raise the question of how excitatory and inhibitory plasticity can cooperate to enable the development of an efficient stimulus code.

Network models have proposed neural-level mechanisms of sparse code formation (Olshausen & Field, 1996) based on Hebbian plasticity. However, these models typically rely on simplified learning dynamics (Savin et al., 2010; Zylberberg et al., 2011; King et al., 2013) or consider plasticity only at a subset of projections in the network (Sadeh et al., 2015; Miconi et al., 2016), not addressing the development of feedback-based gain control. As such, it remains unclear how functional input encoding can emerge during development in a more detailed V1 circuit model.

We here propose how a single underlying mechanism - the influence of inhibitory plasticity on excitatory plasticity - is sufficient to explain both, the observed feed-forward tuning and neuronal gain-control by feedback processing, which we demonstrate in a spiking network model of V1 layer 4. To test for an additional influence of inhibitory strength on the emergence of feed-forward tuning, we varied the balance between excitation and inhibition in the network. Our findings support a role for inhibitory plasticity in the joint development of feed-forward tuning and balanced inhibitory feedback currents. Importantly, this balance leads to the spontaneous emergence of contrast-invariant tuning curves, as an inherent phenomenon of the network and its plasticity dynamics. Our results link both representational efficiency and metabolic efficiency to synaptic plasticity mechanisms.

## 2 Results

To investigate the interaction between excitatory and inhibitory plasticity, we designed a spiking network model of V1-layer 4 consisting of an excitatory and inhibitory population, stimulated with natural image patches (**Fig. 1a**) (see **Network input**). The circuit of our neuronal network implements both feed-forward and feedback inhibition, in agreement with anatomical findings (Isaacson & Scanziani, 2011). Although different kinds of inhibitory neurons have been found in the neocortex (Markram et al., 2004; Priebe & Ferster, 2008), our network contains only one population of inhibitory neurons, as a simplification. The size of the inhibitory population was chosen to match the 4:1 ratio between excitatory and inhibitory neurons found in striate cortex (Beaulieu et al., 1992; Markram et al., 2004; Potjans & Diesmann, 2014). The plasticity of the excitatory synapses follows the voltage-based triplet spike timing-dependent plasticity (STDP) rule proposed by Clopath et al. (2010). The strength of the inhibitory synapses changes according to the symmetric inhibitory STDP rule described by Vogels et al. (2011), which achieves homeostasis by maintaining a constant postsynaptic firing rate (*ρ*).

**Figure 1:**
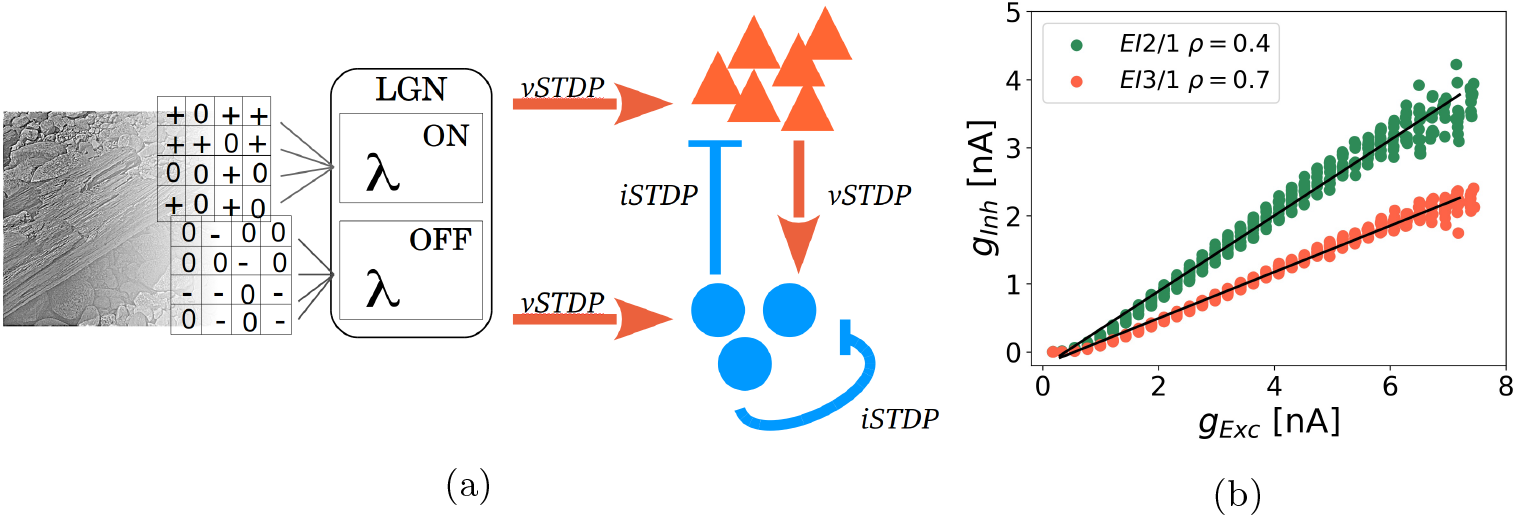
Network with excitatory and inhibitory plasticity rules. (**a**) Whitened image patches of size 12×12 were converted to Poisson spike trains by setting the firing rates of LGN ON- and OFF-populations to the positive and the negative part of the pixel values, respectively. Feed-forward inputs from LGN project both onto excitatory and inhibitory V1 populations, which are mutually connected. The circuit therefore implements both feed-forward and feedback inhibition. Inhibitory interneurons receive additional recurrent inhibitory connections. All excitatory synapses (orange) changes via the voltage-based STDP rule (vSTDP) (Clopath et al., 2010). All inhibitory synapses (blue) changes via the inhibitory STDP rule (iSTDP) (Vogels et al., 2011). Connectivity patterns are all-to-all. Population sizes are: LGN, 288 neurons; V1 excitatory, 144 neurons; V1 inhibitory, 36 neurons. Neurons in the LGN population show Poisson activity and are split into ON- and OFF-subpopulations. (**b**) Inhibitory currents as a function of excitatory currents, averaged across the duration of a stimulus. The post-synaptic target firing rate of the iSTDP rule (*ρ*) controls the excitation to inhibition ratio at excitatory cells. For the *EI*2/1 model (green dots) a value of *ρ* = 0.4 leads to higher inhibitory currents than *ρ* = 0.7 for the *EI*3/1 model.

To analyze the influence of inhibitory plasticity on excitatory plasticity, we used two approaches. First, we investigate how the balance between excitation and inhibition influences the emergence of neuronal gain-control and feed-forward tuning, by comparing a network with a 2: 1 ratio of excitation to inhibition (E/I ratio) to a model version with a 3: 1 E/I ratio, averaged on 10,000 natural scene patches (**Fig. 1b**). This ratio is adjusted via the *ρ* parameter exclusively. Additionally, we blocked inhibitory synapses after learning to investigate the dynamic effects of inhibition on network coding (called *blockInh*). To analyze the influence of inhibition during learning after all, a further model configuration did not contain any inhibitory synapses (called *noInh* model) and learns with the absence of inhibition.

Second, to analyze if plastic inhibition has a measurable effect during learning, we deactivated plasticity selectively at specific connections for two model variations: Only at the inhibitory feedback connections (called *fix fb inh*) and at all excitatory projections to the inhibitory population (called *fix ff inh*). We used shuffled weight matrices from a successfully learned *EI*2/1 model for all connections to ensure that the network will have an E/I ratio comparable to networks where all synapses are plastic. Only the incoming excitatory weights of the excitatory population are chosen anew from a normal distribution. To verify that learning is successful with the shuffled pre-learned weights, we trained one model variation where all connections are plastic(see **Fig. S1**). While we vary the inhibitory influence, the feed-forward synapses to the excitatory population are plastic in all model configurations.

In all model configurations, the populations consist of the same number of neurons and synapses between them. Each model configuration was repeated 20 times. If not mentioned otherwise, initialized with randomly chosen weight values, to test the stability and reproducibility of the observed outcomes.

We first analyze the structural characteristics of the network as a consequence of the learning process, and then present its functional properties. In both cases, we investigate the effect of plastic vs fixed synapses and different inhibitory strengths.

### Emergence of diversely tuned receptive fields

The receptive fields of V1 simple cells are often described by Gabor functions (Jones & Palmer, 1987; Ringach, 2002; Spratling, 2012). We observe the emergence of Gabor-like receptive fields in our network for the excitatory and inhibitory population with the spike triggered average method (STA, see **Receptive field mapping**). Without inhibition, most of the receptive fields have a similar orientation and position (**Fig. 2a**), as it is to be expected from the chosen learning rule, see also Clopath et al. (2010). In contrast, the presence of plastic inhibition during learning resulted in a higher diversity of receptive fields with a more complex structure for the excitatory population (**Fig. 2b**) and the inhibitory population (**Fig. 2c**).

**Figure 2:**
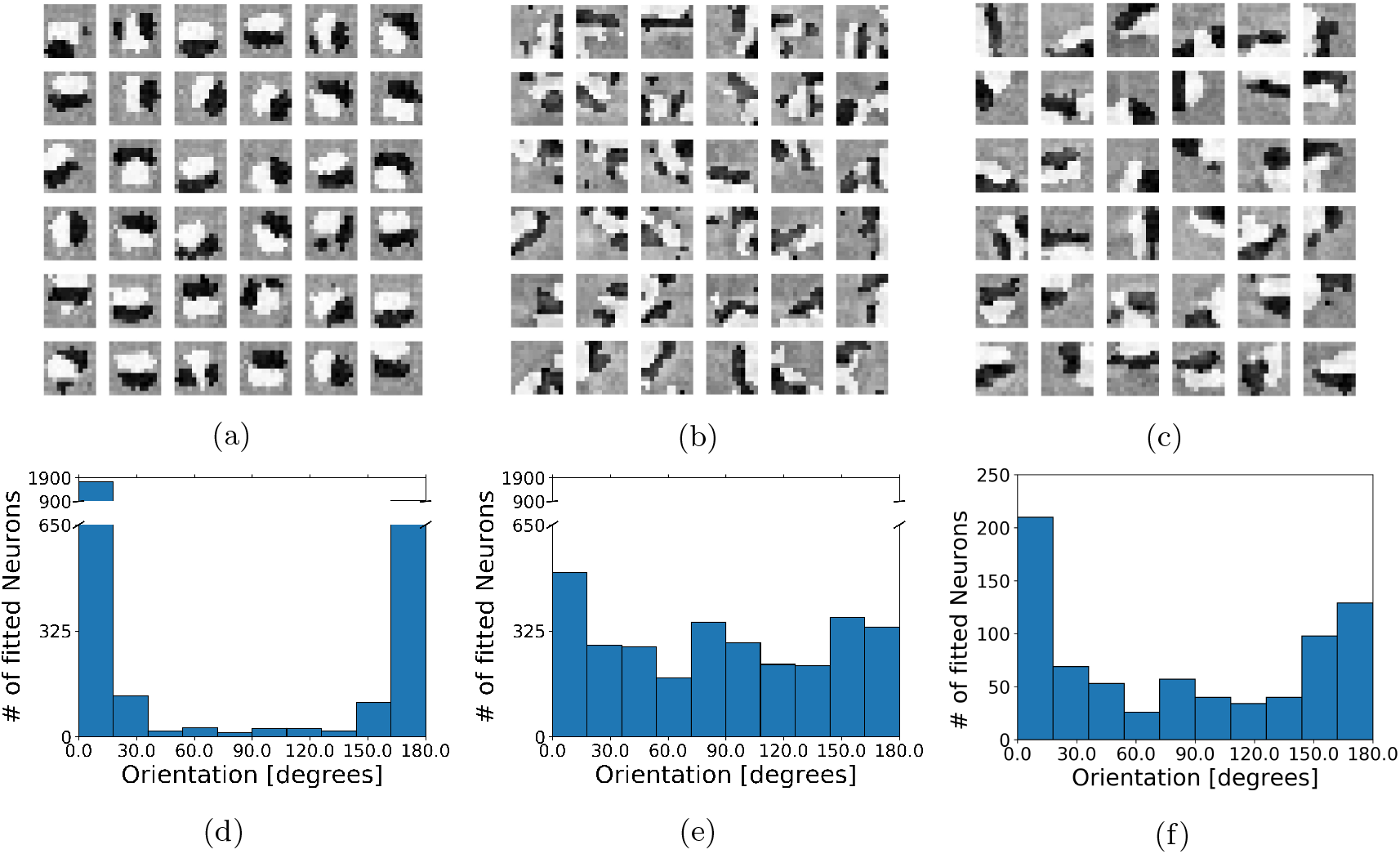
Tuning diversity requires inhibition during learning. (**a - c**) Learned response profile of 36 excitatory neurons from the *noInh* model (**a**), of 36 excitatory neurons from the *EI*2/1 model (**b**), and of all 36 inhibitory neurons from the *EI*2/1 model (**c**), measured with the spike triggered average method. Lighter pixels represent positive values and darker values represent negative values. (**d - f**) Histogram of the spatial orientation across 20 model runs, of the *noInh* model’s excitatory population (**d**), the *EI*2/1 model’s excitatory population (**e**), and the the *EI*2/1 model’s inhibitory population (**f**). Note the strong clustering of orientations in the noInh model (**d**). The spatial orientation are measured by presenting sinus grating on different orientations (see **Tuning curves and orientation selectivity**).

We observed the emergence of stable receptive fields after presenting approx. 200,000 stimuli (see supp. **Fig. S2** and supp. **Fig. S3**). We presented another 200, 000 stimuli to ensure that all synapses reach a stable state. The measured receptive fields showed a strong tendency for weight values to cluster around the minimum or the maximum value (see supp. **Fig. S4**). This is a known characteristic of the learning rule chosen for excitatory synapses, which enforces strong synaptic competition (Clopath et al., 2010; Miconi et al., 2016).

To measure the preferred orientation of each neuron, we presented sinus gratings with different orientations (see **Tuning curves and orientation selectivity**). To quantify the diversity of receptive field orientations across model repetitions, we calculated an orientation diversity index (*ODI*) via the Kullback-Leibler divergence between the measured orientations and an idealized uniform distribution of orientations (see **Eq. 15**). Our calculated *ODI* is the exponential function of the Kullback-Leibler divergence and thus, higher values indicate a more uniform orientation distribution, which means a higher orientation diversity (see **Orientation diversity**).

A broader range of orientations emerged in the networks with inhibition (**Fig. 2e**). Without inhibition, most receptive fields converge to a preferred orientation around 0° or 180° (**Fig. 2d**). In the model with weaker inhibition (*EI*3/1), receptive fields converge to a very similar orientation distribution than in the *EI*2/1 model (see supp. **Fig. S5**). This is mirrored in the orientation distribution (**Fig. 3**). These results suggest that the presence of inhibition is more important for the emergence of receptive field diversity than its strength. In earlier studies of simple cells in the cat visual cortex, a broad distribution of different oriented simple cells has been reported, with a tendency to more cells selective for horizontal stimuli (Rose & Blakemore, 1974), vertical stimuli (Chino et al., 1980) or both (Berman et al., 1987). In our simulations, both models with inhibition (*EI*2/1 and *EI*3/1) show a broad distribution with a slightly higher number of cells with a preference for horizontal and vertical stimuli (see **Fig. 2e** and supp. **Fig. S5**).

**Figure 3:**
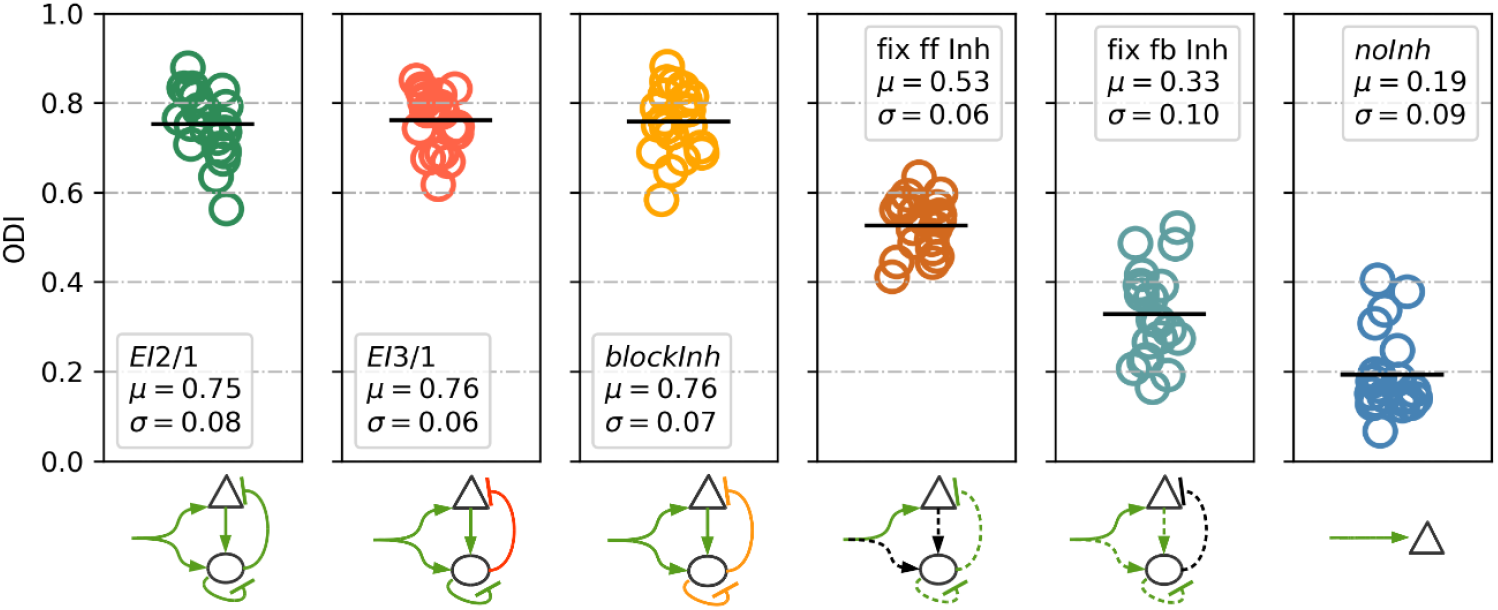
Tuning diversity is improved by plastic feed-forward and feedback inhibition. The orientation diversity index (*ODI*) is calculated via the Kullback-Leibler divergence between an idealized orientation distribution and the measured distribution in the network. The exponential of the divergence value is taken, so that higher values indicate a more uniform orientation distribution. Green arrows indicate plastic synapses, black arrows denote fixed synapses. Orange arrows indicate weaker synaptic connections. Dotted lines indicate that the model weights were initialized with shuffled values from weights of a previous run of the *EI*2/1 model. The highest diversity of RFs is observed for all models with fully plastic inhibition during learning (*EI*2/1, *EI*3/1, and *blockInh* models). Abolishing plasticity at feed-forward inputs to inhibitory neurons led to a moderate decrease of orientation diversity (*fix ff inhmodel*). Blocking plasticity at inhibitory feedback synapses onto excitatory neurons led to a stronger decrease in orientation diversity (*fix fb inhmodel*). The lowest diversity was observed in the *noInh* model, were inhibition was fully absent.

In addition, the inhibitory cells in the *EI*2/1 models also become selectively tuned, with a clear preference at 0° and 180° (**Fig. 2f**), as well as the inhibitory cells in the *EI*3/1 models (see supp. **Fig. S5**). This is in line with recent experimental reports of tuned inhibition in ferret V1 (D. E. Wilson et al., 2017). However, it is still debated whether tuned inhibition is as a general property of the visual system. For example, in mouse V1, recent research has identified inhibitory interneurons which are non-selective for orientation (Kerlin et al., 2010; Hofer et al., 2011; Bock et al., 2011), very broadly tuned interneurons (Liu et al., 2011), and some subtypes of inhibitory interneurons which have a sharp tuning (Runyan et al., 2010).

To further analyze the influence of fixed and plastic feed-forward and feedback inhibition on the resulting orientation diversity, we used the shuffled weight matrices from a *EI*2/1 model to ensure a comparable balance between excitation and inhibition, except for the feed-forward synapses of the excitatory cells, which are newly chosen from a normal distribution. We observed a reduction of tuning diversity in the *fix ff inh* model, in which the excitatory input weights to the inhibitory cells are unspecific and kept fixed (**Fig. 3**). This presumably led to highly homogeneous activity across the interneuron population. A stronger reduction of tuning diversity occurred in the *fix fb inh* model, in which the inhibitory feedback connections were kept fixed. As a consequence, all excitatory neurons received unspecific inhibitory feedback. As expected, the *noInh* model showed the lowest degree of tuning diversity in the absence of any inhibition.

### Emergence of structured feed-forward and recurrent connectivity

As both, the excitatory and inhibitory cells in our network developed a tuning for orientation and position, we expected that their modifiable synaptic connections developed a specific pattern reflecting activity correlations (King et al., 2013; Sadeh et al., 2015). For an exemplary model simulation, our analysis confirmed that excitatory neurons developed strong connections to inhibitory neurons with similar orientation tuning (**Fig. 4a**, top). Inhibitory weights to the excitatory layer showed a similar pattern, although with somewhat reduced specificity (**Fig. 4a**, bottom). This implements an indirect inter-neuron connection between two excitatory neurons via mutually connected inhibitory neurons, to inhibit each other maximally. The development of recurrent inhibitory synapses between similarly tuned inhibitory cells can be observed as well (**Fig. 4b**). We next analyzed the connectivity structure based on all model repetitions as follows: First, for any pair of neurons sharing a synaptic connection, we calculated the template match between their receptive fields. Second, we binned the weight values and template match values for all neuron pairs from all model repetitions. Finally, we plotted the average weight strength as a function of the average template match for all neuron pairs per bin (**Fig. 4c**). For both models with plastic inhibition (the *EI*2/1 and the *EI*3/1 model), we observe that neurons with a more similar receptive field have a higher mutual synaptic weight value. These results are in agreement with recent experimental reports from mouse visual cortex (Cossell et al., 2015).

**Figure 4:**
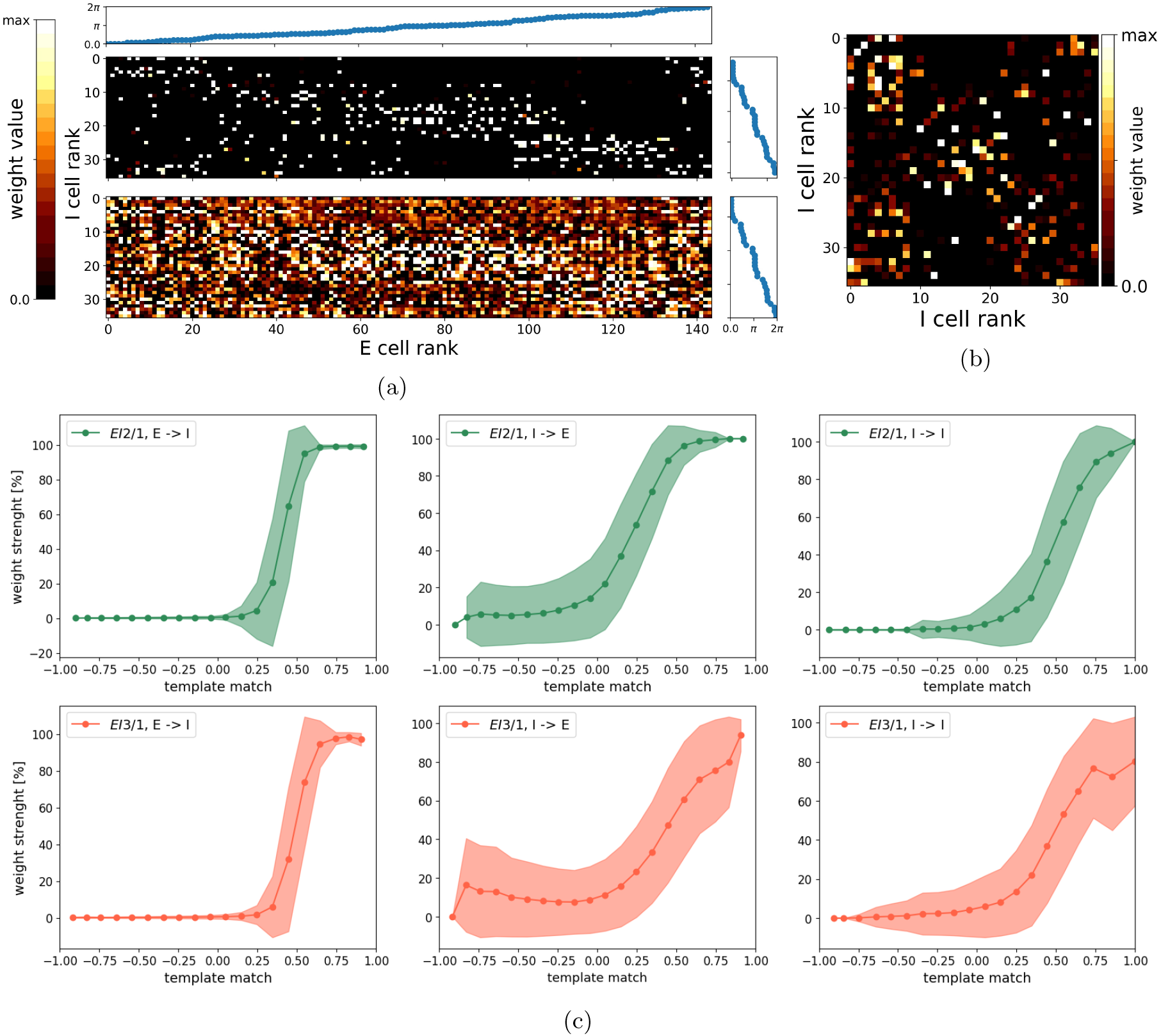
Synaptic connections reflect tuning similarity. Weight matrices from the excitatory to the inhibitory population (and vice versa) (**a**), sorted by the receptive field orientation, and for the lateral inhibitory weights (**b**). **a**,Top: Weights from the excitatory to the inhibitory population. **a**, Bottom: Weights from the inhibitory to the excitatory population. For display, all weight matrices were normalized by the maximum value. All weights are from the *EI*2/1 model. (**c**) Normalized synaptic strength as a function of the template match between the pre- and postsynaptic neuron’s receptive fields for the *EI*2/1 (first row) and the *EI*3/1 (second row) model. Shaded areas denote the mean +/- standard deviation. As expected, we observed strong weights between neurons with highly similar receptive fields, and near-zero weights between neurons with highly dissimilar receptive fields. For neurons with a moderate degree of RF similarity, we observed a steep transition from weak to strong weights at the E-I projection. At the I-E and I-I projections, this transition was more gradual.

### Inhibition controls response decorrelation

We observed that the different levels of inhibition in the *EI*2/1 and *EI*3/1 models led to similar orientation distributions. To investigate if response correlations between neurons only depend on the orientation similarity or whether lateral inhibition has an additional decorrelation effect (as mentioned in previous modeling approaches of Wiltschut & Hamker (2009); Savin et al. (2010); Zylberberg et al. (2011); King et al. (2013)), we analyzed the development of correlations during receptive field learning (**Fig. 5a**). During the first 250,000 of all 400,000 input stimuli, a weak reduction of the correlation can be observed in the *noInh* model. The *EI*2/1 model showed a pronounced decrease of correlations across learning, with the highest reduction occurring in the early phase of learning showing the highest amount of changes of the feed-forward weights. Weaker feedback inhibition (*EI*3/1 model) led to weaker decorrelation of neuronal activity.

**Figure 5:**
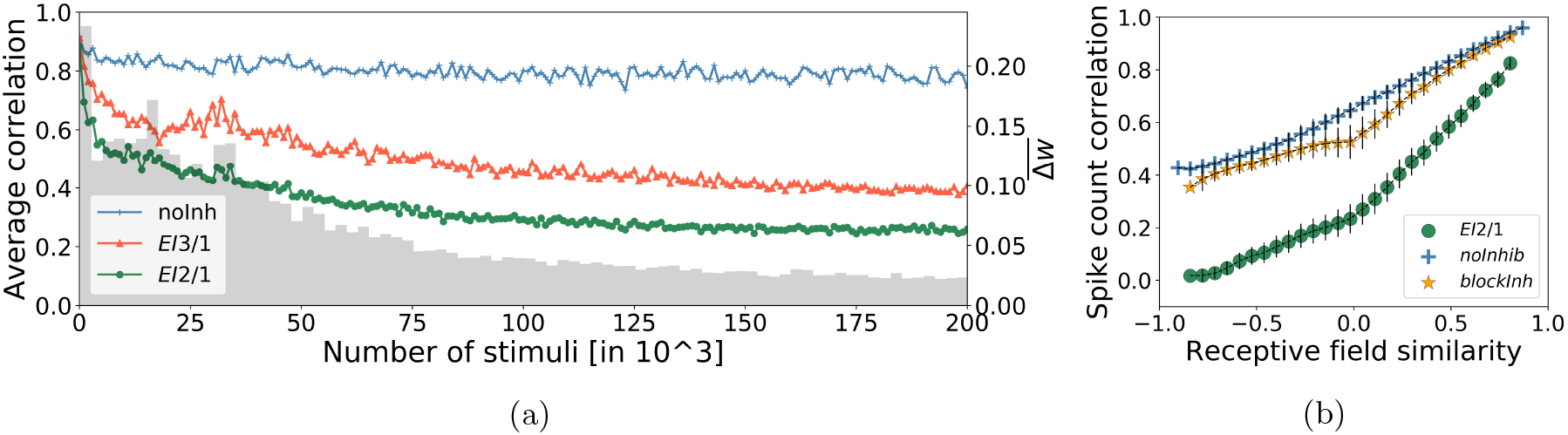
Inhibitory strength influences the response decorrelation. (**a**) The development of mean response correlation and weight change at the LGN excitatory synapses across learning. Stronger inhibition, in the *EI*2/1 model, leads to a stronger decorrelation of the neuron responses during learning (compare green with red (*EI*3/1) line). Mean response correlation changed only very slightly without inhibition (blue line). The change in the synaptic weights (gray bars) decreases over the developmental process, indicating the emergence of stable receptive fields. (**b**) Response correlation is higher for neurons with more similar receptive fields. Blocking inhibition (yellow line) after learning reveals that inhibition leads to a overall decrease of the response correlation (green line).

Smith & Kohn (2008) recorded the neuronal activity in V1 of macaque monkeys during the presentation of drifting sinusoidal gratings and reported a dependence of pairwise response correlations on orientation tuning similarity. We performed a similar analysis of our model data, to analyze the effect of feedback inhibition on the response correlation with respect to the orientation selectivity. We sorted all cell pairs by similarity, grouped them into 30 equally-spaced bins, and averaged their response correlation values within each bin, based on natural scene stimuli (details see **Methods**) (**Fig. 5b**). In both models without inhibition, we observed a mean response correlation of ≈ 0.95 for cell pairs with highly similar receptive fields. With inhibition, this value dropped to ≈ 0.8. By contrast, cell pairs with dissimilar receptive fields showed average correlation values of around 0.4 for the *noInh* and the *blockInh* model. Here, inhibitory processing substantially reduced the mean correlation to near zero-values for the *EI*2/1 model. A comparison between the *EI*2/1 model and its counterpart with blocked inhibition shows that dissimilarly tuned neuron pairs are more strongly decorrelated than pairs with highly similar tuning. At a first glance, this pattern contrasts with the emergent connectivity structure: The connectivity pattern favors strong mutual inhibitory connections between inhibitory neurons which receive projections from (and project back to) excitatory neurons with similar tuning, creating strong reciprocal inhibition (**Fig. 4a** and **Fig. 4b**). However, our observation of target-specific decorrelation is best understood by considering that correlated spike counts can arise both through a similarity of tuning and through unspecific baseline activity, caused by contrast differences. Natural image patches are likely to evoke broad excitation among many cells, leading to different neuronal responses as sinusoidal gratings (Kayser et al., 2003). Due this, studies measuring the pairwise response correlation with sinusoidal gratings, reported a stronger decorrelation effect between similar neurons (Smith & Kohn, 2008; Denman & Contreras, 2013). Despite that, studies presenting natural scene inputs to measure the neuronal response correlation reported higher correlation values in comparison to sinusoidal gratings (Froudarakis et al., 2014; Martin & Schröder, 2013) or more similar values as reported here (Weliky et al., 2003). The correlation between dissimilarly tuned neurons is most likely caused by the activity baseline, which is strongly reduced by inhibition. Besides, similarly tuned cells will retain strongly overlapping tuning curves even after reduction of unspecific activity, associated with strong correlation of their mean response (Averbeck et al., 2006). Our observation that blocking the inhibitory processing leads to an overall increase of activity correlation is in line with previous studies. Sippy & Yuste (2013) reported an increase of activity correlation between principal cells from 0.31 up to 0.66 by reducing inhibition pharmacologically in thalamocortical slices from mice (without considering receptive field similarities). A similar increase is observable if the compare the mean pairwise correlation from the *EI*2/1 model (0.32) and the *blockInh* counterpart (0.60).

### Inhibitory feedback shapes tuning curves

To quantify the effect of inhibition on the magnitude of individual neuronal responses, we measured orientation tuning curves of each neuron by sinusoidal gratings. For all approaches and model variants, the maximum firing rate in the input was set to ≈ 85*Hz* to obtain sufficiently high activity levels. We observed high baseline and peak activity in both model variants without inhibition (**Fig. 6a**). However, activity levels in the *blockInh* model were lower than in the *noInh* model, likely owing to its smaller and more dispersed receptive fields. As expected, the model with active inhibitory feedback showed the lowest firing rate to input ratio. To obtain a measure of tuning sharpness, we next estimated the orientation bandwidth (OBW) of the excitatory population, based on the measured tuning curves. As expected, and consistent with previous observations (Isaacson & Scanziani, 2011; Stringer et al., 2016), our model shows a sharpening effect through inhibition (**Fig. 6b**).

**Figure 6:**
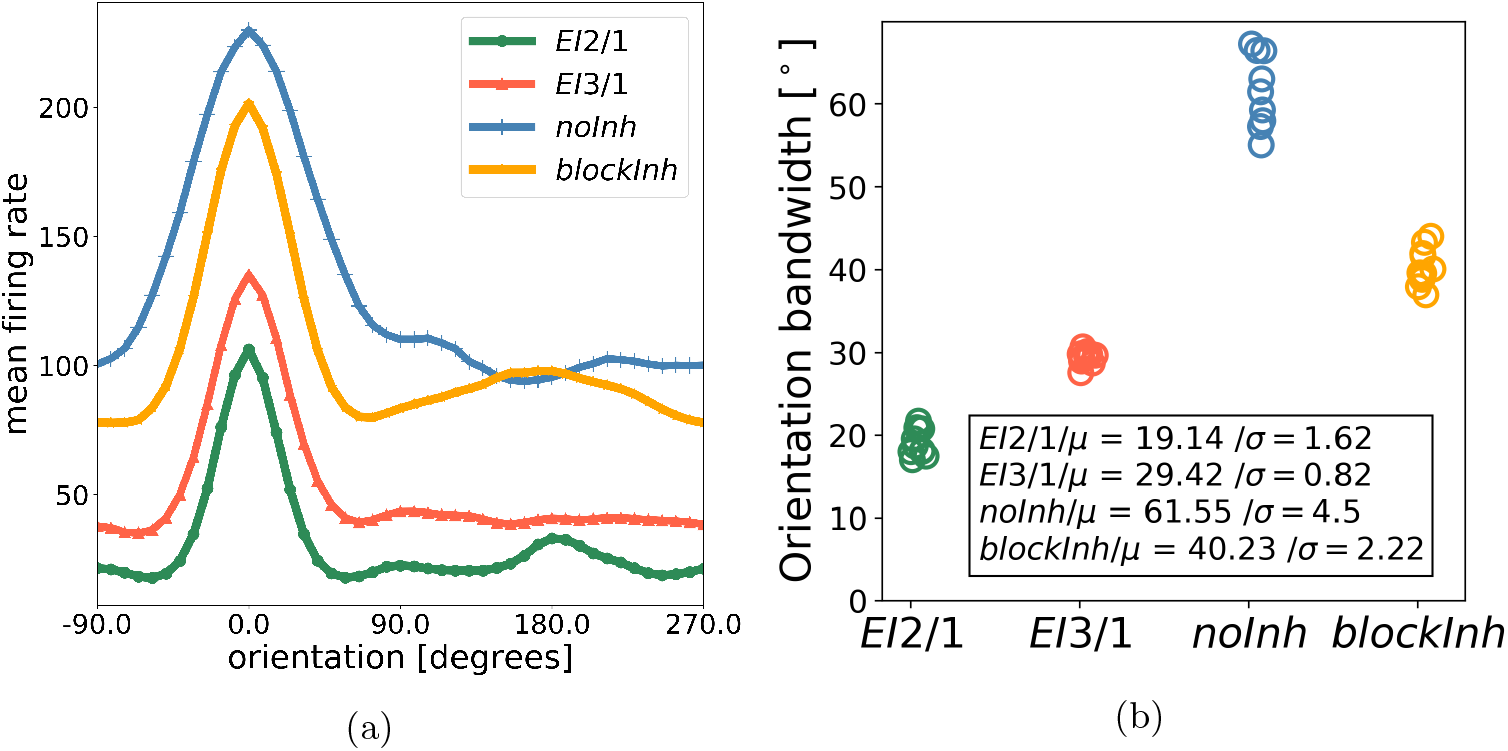
Inhibition controls tuning curve sharpening. (**a**) Average tuning curve of all excitatory cells in the *EI*2/1 model, the corresponding counterpart with blocked inhibition, and the no inhibition model. (**b**) The orientation bandwidth (OBW) of cells in all three models. Every point represents the average OBW resulting from model simulation. Smaller OBW values correspond to narrower tuning curves. As expected, the *EI*2/1 model (green) shows the narrowest tuning curves. The slightly reduced inibitory strength in the *EI*3/1 model (red) leads to moderately broader tuning curves. Fully blocking inhibition post-learning leads to both wider tuning curves and increased baseline activity in the *blockInh* model (yellow). The broadest tuning curves and highest baseline activity were observed in the *noInh* model (blue), which produced relatively large receptive fields.

Duo to the same overall magnitude of inhibitory feedback as for the *EI*2/1 model, we assume for the *fix ff inh* and the *fix fb inh* a highly similar behavior, as it has been reported in previous work that broad or untuned inhibition causes tuning sharpening (Ben-Yishai et al., 1995; Troyer et al., 1998; Priebe & Ferster, 2008) (data not shown).

### Spontaneous emergence of contrast-invariant tuning curves

Besides the sharpening of tuning curves, previous models suggest a role of inhibition in the invariance to input contrast changes (Troyer et al., 1998; Ferster & Miller, 2000; Priebe & Ferster, 2008). However, those models assume hard-wired connectivity, and propose push-pull or anti-phase inhibition (Troyer et al., 1998; Ferster & Miller, 2000). Contrast-invariant V1 simple cells have been found in different mammals such as, cats (Skottun et al., 1987; Finn et al., 2007) or ferrets (Alitto & Usrey, 2004), based on sinusoidal gratings with different contrast strength. We use the same approach (see **Tuning curves and orientation selectivity**) to measure the tuning curves and calculated the averaged OBW over all excitatory cells for the different contrast levels (**Fig. 7a**). Interestingly, the OBW is constant only for the *EI*2/1 model. For the model with weaker inhibition (*EI*3/1 model) and the model without inhibition (*noInh*), the OBW increases for higher input contrast values. Similarly, we observed a contrast-dependent increase in tuning width when inhibition was blocked after learning (*blockInh*). As it has been shown that random feedback inhibition is sufficient for the emergence of contrast invariant tuning curves Ben-Yishai et al. (1995), we omit data for the *fix ff inh* and *fix fb inh* models for clarity of display.

**Figure 7:**
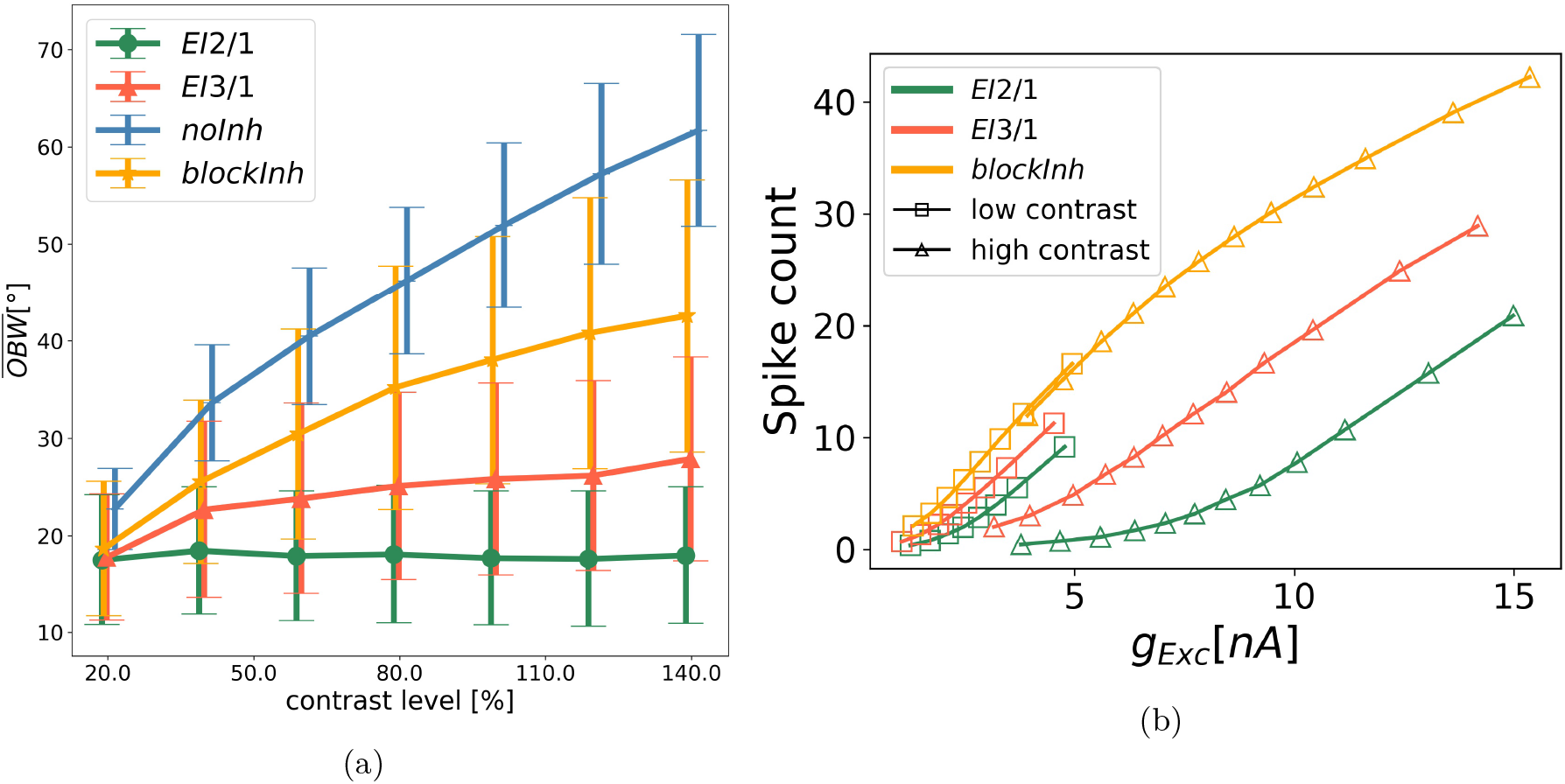
Response gain control by inhibition. (**a**) Mean OBW as a function of the contrast level in the input. Whiskers represent the standard deviation. Data from the *EI*2/1 model (green line), the model with all synapsed are from and to the inhibitory population are random and fixed (gray line), *EI*3/1 model (red line), and *noInh* model (blue line). (**b**) Spike count as a function of the excitatory input current for the *EI*2/1 model (green line), the *EI*3/1 model (red line) and the *blockInh* model (orange line). Data are taken from the sinusoidal tuning curve measurement, sorted by input current. Squares: Low input contrast. Triangles: High input contrast. Contrast-invariant tuning is only present in the *EI*2/1 model, while all other models show varying degrees of contrast-dependent widening of tuning curves.

To understand how the strength of inhibition affects contrast tuning curves, we compared the *EI*2/1 with the *EI*3/1 model with regard to their spike count, average membrane potential, and the average of the summed synaptic input current, for different contrast levels. At any contrast level, the activity of neurons in the *EI*2/1 model remains strongly suppressed at non-preferred orientations and increases around the preferred orientation (**Fig. 8a**). By contrast, the *EI*3/1 model shows increased activity for high input contrast at all orientations (**Fig. 8b**). This results in increased OBW values for higher input contrast (see also **Fig. S9 for normalized spike counts**). Interestingly, for the non-preferred orientation, the average membrane potential the *EI*2/1 model is less hyperpolarized for lower contrast than for higher contrast. For higher contrast, the average membrane potential increases at the preferred orientation and is substantially stronger than for lower contrast. Both curves intersect around − 50*mV*, close to the resting state spiking threshold (−50.4*mV*) (**Fig. 8c**). This can be explained with the average input current: At higher contrast levels and non-preferred orientations, the feedback inhibitory current increases more strongly than the excitatory current and nearly compensates it (**Fig. 8e** and **S3 a**), providing hyperpolarization of the membrane potential. This compensation of excitation decreases around the preferred stimulus, where the membrane potential exceeds the spiking threshold. In comparison, the membrane potential for the *EI*3/1 model increases proportionally with the total input current caused by higher input contrast (**Fig. 8d**, **Fig. 8f** and **S3 b**). This suggests that the contrast-invariant tuning of the *EI*2/1 model depends on an appropriate balance between excitation and inhibition.

Based on the observation of contrast invariant tuning curves, we conclude that feedback inhibition modulates the neuronal gain controlled by input orientation and contrast. **Fig. 7b** shows the average response gain for the excitatory population, averaged across the whole population, and sorted by the input current(see **Neuronal gain curves** for more details). We show the response gain curves for low and high contrast stimuli. For the model with blocked inhibition (*blockInh*), the gain curve is unaffected by contrast and follows the activation function defined by the neuron model. The firing rate to input ratios of neurons in the *EI*2/1 model are strongly reduced relative to the *blockInh* model, but this gain modulation is contrast-dependent, as the highest reduction of firing rates is observed for high contrast. This shows that the effect of inhibition on the neuronal gain function not only depends on the amount of excitatory input, but also on the stimulus orientation and contrast strength.

**Figure 8:**
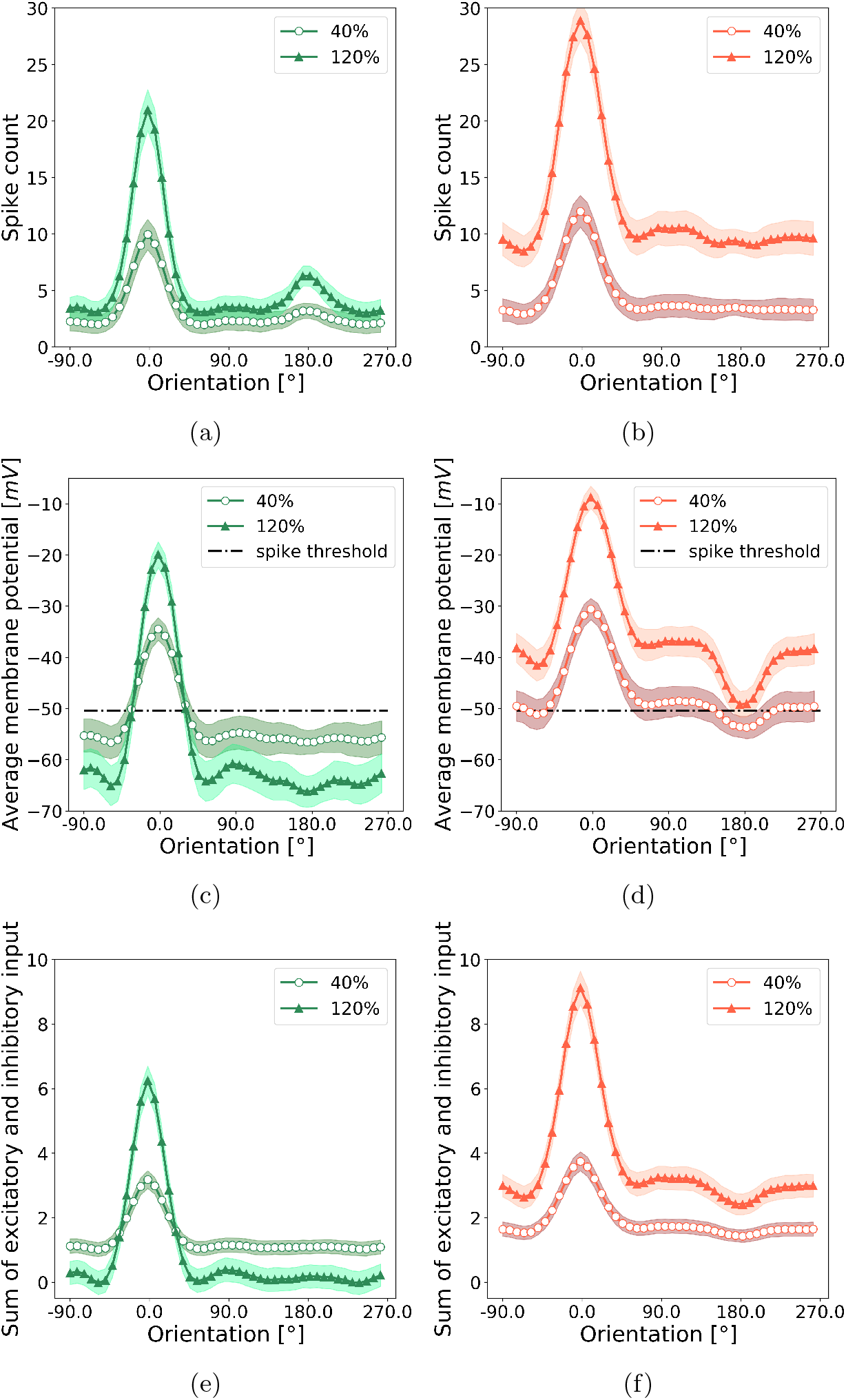
Emergence of contrast-invariant responses. (**a**) Average neural tuning curves for low and high contrast stimuli in the *EI*2/1 model, (**b**) and the *EI*3/1 model. (**c**) Average membrane potential (averaged across all neurons in the excitatory population) as a function of orientation and contrast level for the *EI*2/1 model, (**d**) and the *EI*3/1 model. (**e**) Sum of the excitatory and inhibitory input currents as a function of orientation and contrast level for the *EI*2/1 model, (**f**) and the *EI*3/1 model. In the *EI*3/1 model, high-contrast stimuli with non-preferred orientations are associated with very different dynamics than in the *EI*2/1 model: In the *EI*2/1 model, the sum of excitatory and inhibitory currents is near zero for non-preferred orientations at high contrast (e). In the *EI*3/1 model, the total synaptic current (f) remains large enough to elicit considerable membrane depolarization for non-preferred orientations at high contrast (d), reflected in elevated baseline activity and broader tuning (b).

### Sparseness is increased by both, inhibition and tuning diversity

As we observed that inhibitory processing led to an increase in the selectivity to artificial stimuli, we asked whether inhibition contributed to a sparser population code for natural images. We first compared the overall spiking behavior based on raster plots of network responses to five example image patches, for the *EI*2/1 (**Fig. 9a**) and the *blockInh* model (**Fig. 9c**). The model with active inhibition showed sparser firing and a less synchronous spiking behavior than the model with blocked inhibition. Second, to quantify this effect, we measured the population sparseness for all model configurations, based on the responses to 10.000 natural image patches (**Fig. 9b**). The highest sparseness value (0.62) was observed in the *EI*2/1 model, 0.54 for the *blockInh* model and the lowest sparseness value (0.43) in the *noInh* model. Interestingly, the development of a higher diversity of receptive fields had a stronger influence on the population sparseness than inhibitory processing: Sparseness values differed more strongly between the model configurations without inhibition, the *noInh* and *blockInh* model, than between the *EI*2/1 and its blocked counterpart, which share the same feed-forward receptive fields.

**Figure 9:**
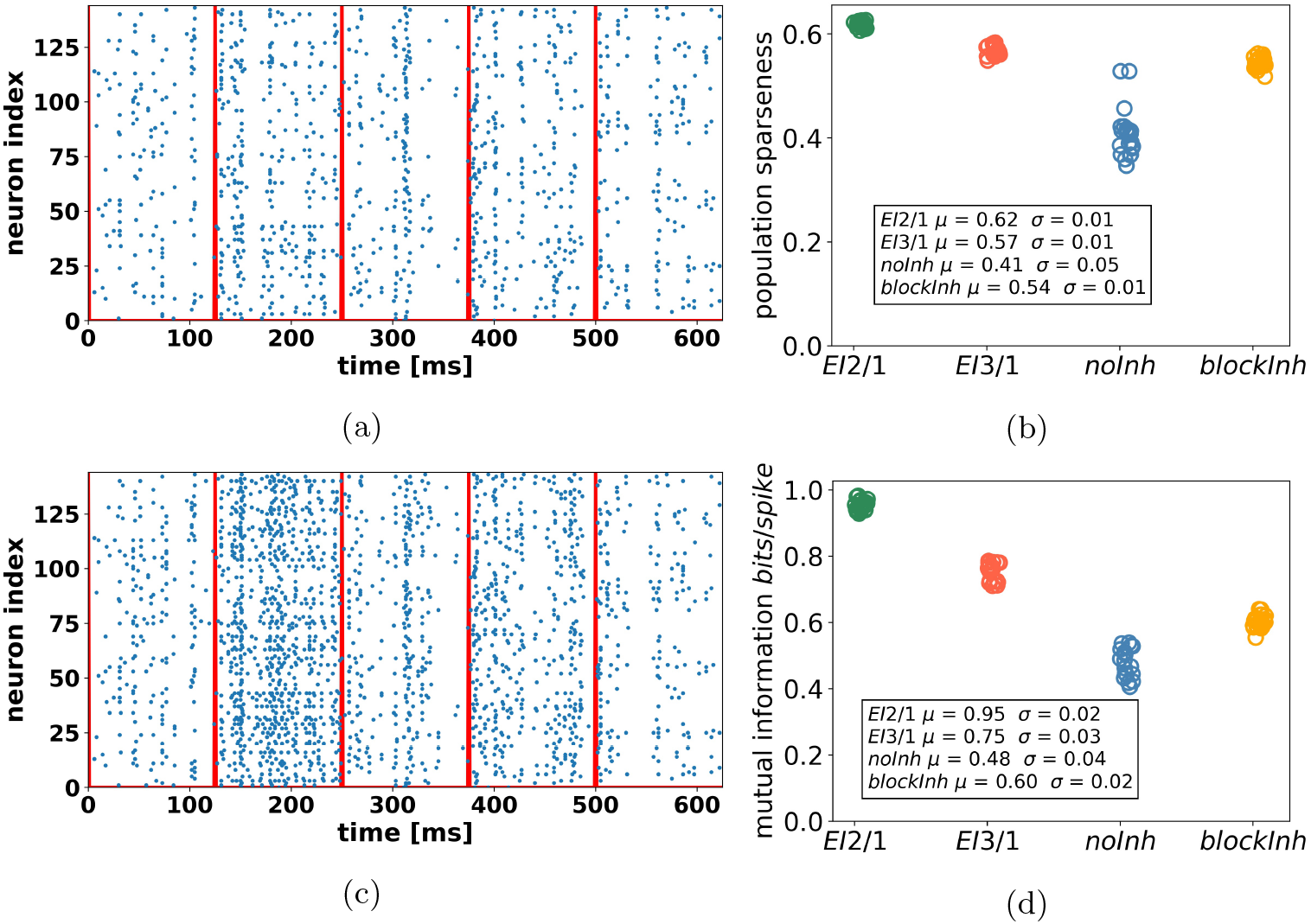
Sparse and efficient input representations through inhibitory processing. (**a**) Raster plot of the excitatory population for the *EI*2/1 model, same for the *blockInh* model (**c**). Spikes are recorded on the same five natural image patches. The red lines show the stimulus onset. (**b**) Population sparseness for the *EI*2/1, the *blockInh*, and the *noInh* model, averaged across 10.000 natural scene patches. Higher value represent a higher sparseness of population activity. (**d**) Mutual information in *bits/spike* for the same three models as in (**b**). (**b**) and (**d**) show data from 20 independent simulations per model configuration. Note the more synchronous population activity in the *noInh* model (c), associated with reduced sparseness (b) and lower information content (d). While blocking inhibition post-learning in the *blockInh* model decreases sparseness only moderately, it considerably reduces the information per spike.

### Metabolic efficiency benefits from strong feedback inhibition

The efficiency of information transmission (such as the numbers of spikes to represent specific input stimuli and the amount of information transmitted via a spike), or metabolic efficiency, is associated with the observed increase of the population sparseness (Spanne & Jorntell, 2015). To quantify the metabolic efficiency, we calculated the mutual information between input and response (**Sec. Mutual information**). This analysis revealed a strong impact of inhibition on transmission efficiency (**Fig. 9d**), normalized by spike count. The *EI*2/1 model shows the highest amount of information per spike (0.96 *bits/spike*). A lower inhibition strength in the *EI*3/1 model leads to a lower transmission efficiency (0.77 *bits/spike*). Both models without inhibition were associated with the least efficient population coding, with a lower value for the *blockInh* model, caused by a more diverse receptive field structure. To analyze further how the increase in information transmission was achieved, we calculated the discriminability index *d′* on 500 randomly chosen natural scene patches to quantify the trial-to-trial fluctuation. We observed that higher *d′* values were associated to both, high tuning diversity and the presence of inhibition(see supp **Fig. S6**). The improvement in discriminability is likely caused by a reduction of unspecific activity by inhibition, associated with more reliable stimulus representations, as observed in cat V1 (Haider et al., 2010) and mouse V1 (W. Zhu et al., 2009). In summary, our results show that the inhibitory processes in our models suppress redundant spikes which convey little information about the current stimulus (Kremkow et al., 2016).

Metabolic efficiency has also previously been linked to a minimum wiring principle (Graham & Field, 2010) between neurons or cortical areas (Graham & Field, 2010; Mitchison & Barlow, 1991). While it would be interesting to explore effects of structural plasticity on metabolic efficiency, we here focused on the effects of inhibition.

### Input reconstruction benefits from plastic inhibition

We assume that a diversity of receptive fields, which encode the relevant input features, is crucial to provide an input representation without loss. To measure the quality of the input representation and to compare our model with existing sparse coding models, in terms of stimulus encoding, we calculated the image reconstruction error (IRE), which measures the mean-square error between the input image and its reconstruction obtained by linear decoding (see **Image reconstruction error**). We plot the IRE as a function of the receptive field diversity, measured by the orientation diversity index (*ODI*) as described previously (see **Orientation diversity**). The *EI*2/1 model with active and plastic inhibition during learning showed the lowest reconstruction error value (0.74), with a high ODI value (0.75) (**Fig. 10**). By contrast, we observed a substantially smaller ODI value if there is no inhibition (*noInh* model) at all during learning (0.19), resulting in a higher reconstruction error (1.06). When the inhibition was blocked in the *EI*2/1 model after learning, the IRE shows a slight increase to a value of 0.79 (*blockInh* model). For the *EI*3/1 model we observed a similar IRE of 0.75 and a similar ODI value (0.76), indicating that the strength of inhibition during learning did not influence the emergence of receptive field diversity nor the input encoding quality.

**Figure 10:**
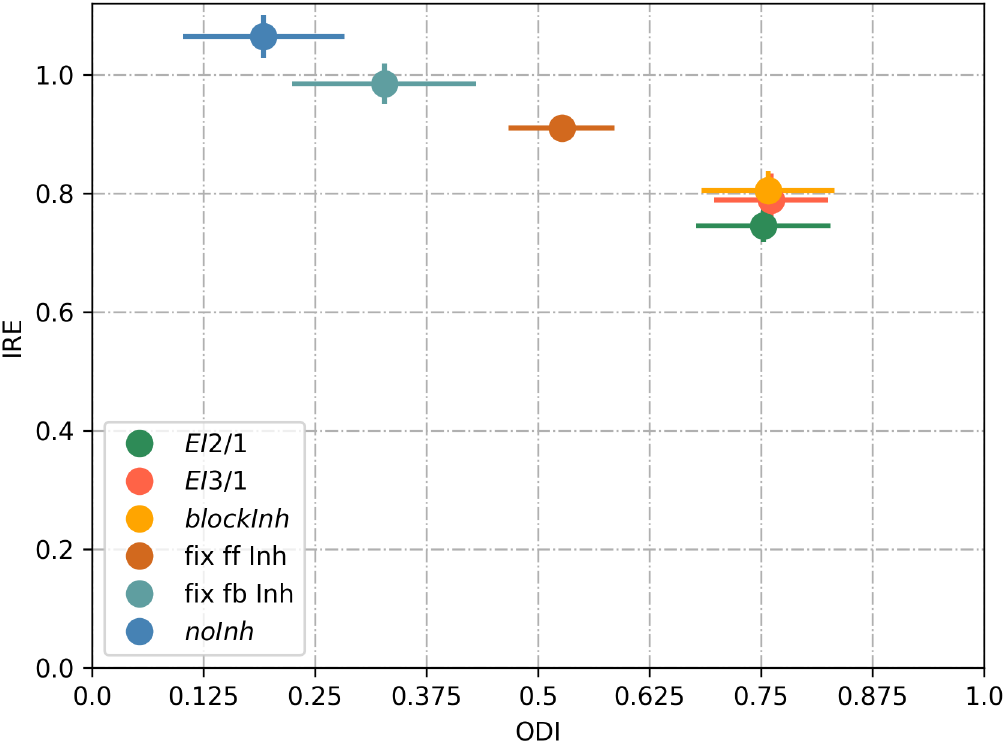
Plastic inhibition during learning improves input encoding quality via higher orientation diversity. Image reconstruction error (IRE) as a function of the orientation diversity index (ODI), for the *EI*2/1 model (green dot),the *EI*3/1 model (red dot), the *blockInh* model (orange dot), model with fixed feed-forward inhibition (brown dot), model with fixed feedback inhibition (light blue dot), and the *noInh* model (dark blue dots). IRE is calculated as the mean-square error between input image and the reconstruction. A better reconstruction is represented by smaller values for the IRE and is associated with a higher orientation diversity (presented by higher ODI values). Data shown from 20 independent simulations per model configuration.

If the feed-forward input to the inhibitory population is random and fixed during learning (*fix ff inh* model), the receptive fields of the excitatory population are less diverse, and the reconstruction error increases (0.91). A fixed inhibitory connection to the excitatory population (*fix fb inh* model) leads to a slightly higher reconstruction error (0.97) and a less diverse receptive field orientations (ODI of 0.33). This demonstrates that the plasticity of both the inhibitory feedback connections and the excitatory feedforward connections to the inhibitory population leads to a better input representation, as a consequence of a higher receptive field diversity. Using fixed inhibitory feed-forward and feedback connections lead to a similar result then having only fixed feedback inhibitory connections (see **Fig. S1**)

To verify that the influence of plastic inhibition is the cause for a more receptive field diversity, and not a mechanisms of the chosen excitatory learning rule, we replaced the Clopath et al. (2010) learning rule with the triplet STDP learning rule from Pfister & Gerstner (2006). We add a spike-traced based homeostatic mechanism (as suggest in Pfister & Gerstner (2006)) to realize receptive field learning and tested fixed feed-forward, fixed feedback and non-plastic inhibition in the same way as for our original model. We observed the same reduction of orientation diversity with an increase in the IRE by fixed feed-forward and/or feedback inhibition (see supp. **Fig. S14**). Together, these results indicate that the diversity of receptive fields contributes to the average reconstruction accuracy. Further, after learning, the effect of active inhibition on the encoding quality is negligible. This is important, as inhibition is essential for receptive field diversity, but it may contribute to a loss of information if the neural code becomes too sparse by the suppression of too many feature-coding neurons (Wiltschut & Hamker, 2009).

This is crucial for a robust input representation, where a very sparse representation (or local code) is less robust against noise (Spanne & Jorntell, 2015). We already showed in two previous studies with similar neural networks, how inhibition can increase the robustness against the loss of information in the input (what can be understood as noise) (Kermani Kolankeh et al., 2015; Larisch et al., 2018). Additional, we measured the resulting image reconstruction error with white noise added on a natural scene and observe a higher robustness against noise in models with plastic inhibition (see **Fig. S15**).

## 3 Discussion

Our model suggests that a single underlying mechanism - the interaction of excitatory and inhibitory plasticity - can explain the stable emergence of reliable and efficient input encoding. We have shown that in particular, the combination of plastic inhibitory feedback and plastic feed-forward inhibition has an influence on shaping the receptive fields. Our simulation results are supported by recent physiological findings that inhibitory plasticity influences the mode of operation of excitatory neurons (for example the excitability) (Griffen & Maffei, 2014; Wang & Maffei, 2014; Khan et al., 2018; Znamenskiy et al., 2018), or influences the occurrence of LTP and LTD (Paille et al., 2013; Griffen & Maffei, 2014; Mongillo & Loewenstein, 2018).

Previous models based on STDP rules, which have demonstrated the emergence of V1 simple cells, made several simplifications in terms of the learning dynamics (Savin et al., 2010; Zylberberg et al., 2011; King et al., 2013), or considered plasticity only for a subset of projections (Sadeh et al., 2015; Miconi et al., 2016). These assumptions make it difficult to investigate the influence of plastic feed-forward and feedback inhibition on network dynamics and input encoding. For example, the observation of response decorrelation is a direct consequence of the chosen learning mechanism (Zylberberg et al., 2011; King et al., 2013). Other learning rules have been designed to optimize the mutual information between input and output (Savin et al., 2010). This suggests that a more detailed model of V1 circuit development is necessary to understand the dynamics between excitation and inhibition during learning.To advance our understanding of this process, we investigated a spiking network model of V1 simple cell development, based on two phenomenological learning rules implemented at all synaptic projections.

### Feed-forward and feedback inhibitory plasticity improves orientation diversity and representational efficiency

Our results show that plastic inhibitory feedback as well as plastic feed-forward inhibition influence the development of V1 simple cells, lead to a higher orientation diversity, and improve representational efficiency. Inhibitory plasticity has been reported in numerous physiological studies (Froemke et al., 2007; Carvalho & Buonomano, 2009; Kullmann et al., 2012; Wang & Maffei, 2014; D’Amour & Froemke, 2015; Khan et al., 2018). Previous model studies suggest a role for inhibitory plasticity in controlling the balance between excitation and inhibition (Vogels et al., 2011; Litwin-Kumar & Doiron, 2014), or in enabling stability in recurrent networks (Litwin-Kumar & Doiron, 2014; Sprekeler, 2017). However, there is ongoing discussion about the necessity and role of inhibitory plasticity during learning a functional sensory code book (Griffen & Maffei, 2014; Srinivasa & Jiang, 2013; Sprekeler, 2017), and this issue has received limited attention in model studies so far.

In a model based on a combination of STDP and inhibitory STDP learning rules, Litwin-Kumar & Doiron (2014) showed that inhibitory plasticity is necessary for stable learning in a network with recurrent excitatory connections. Their study used a generic cortical network receiving non-plastic input from a set of 20 artificially stimuli, which in turn resulted in the formation of 20 assemblies representing the input stimuli. They emphasized that inhibitory plasticity acted to equilibrate firing rates in the network, such that different assemblies (each coding for one stimulus) received different amounts of inhibition, preventing dominant activity of single assemblies. Our results of a feature-specific strength of inhibition generalize their finding of firing rate heterogeneity induced by iSTDP from an “assembly code”, in which different stimuli rarely overlap, to the quasi-continuous space of natural visual stimuli. This supports the necessity of the interaction of inhibitory and excitatory plasticity during the development of the visual cortex.

### Emergence of a self-organized balance of excitation and inhibition

Based on natural scene stimuli, we observed in our model that the inhibitory input current to a neuron is proportional to the excitatory input, when the currents are averaged across the duration of a stimulus. However, as we did not observe an equal strength between these currents, excitation is dominant in our network. This indicates a detailed and loose balance (for definition see, Hennequin et al. (2017)) between excitation and inhibition in our network. While a detailed balance has been reported in rat auditory cortex (Dorrn et al., 2010), it is still under discussion if a more loose or tight balance exists in the primary visual cortex of higher mammals (Froemke, 2015). Recent model studies suggest a tight balance between inhibition and excitation (Denève & Machens, 2016) or rather an inhibitory dominated network for stable learning in a network with recurrent excitatory synapses (Litwin-Kumar & Doiron, 2014; Sadeh et al., 2015; Miconi et al., 2016). However, most of these models investigate excitation-inhibition balance in a singe-neuron setup (Deneve & Machens, 2016), or set a subset of synaptic connections fixed (Litwin-Kumar & Doiron, 2014; Sadeh et al., 2015; Miconi et al., 2016). Interestingly, we observed that the ratio between excitation and inhibition changes in our network for different contrast levels of sinusoidal grating stimuli, up to a 1: 1 balance for the highest contrast level for the *EI*2/1 model. This shows that the balance between excitation and inhibition is input-specific.

### Inhibition implements a gain control mechanism and shapes tuning curves

Previous physiological studies found that parvalbumin-expressing (PV) interneurons have a divisive impact on the gain function of pyramidal neurons in the visual cortex, to implement a contrast gain control mechanism (Atallah et al., 2012; N. R. Wilson et al., 2012; Y. Zhu et al., 2015). In our model we observed that the ratio between excitatory and inhibitory currents influences the response of the neuron towards its input. Consequently, feedback inhibition implements a gain control mechanism for the excitatory neurons.

Savin et al. (2010) proposed a rapid intrinsic plasticity mechanism to adapt the neuronal gain function to optimize the information transmission between input stimuli and neuronal output. They suggested that the emergence of V1 simple cell receptive fields depends on the interplay between the adaption of the neuronal gain function and the synaptic plasticity (Savin et al., 2010). By contrast, in our network, changes in neuronal gain curves are caused by feedback inhibition, which adapts at the fast time scale of synaptic plasticity to maintain a given target rate.

In our model, when blocking inhibition after learning, we observed an increase not only in the baseline activity, but also in the orientation bandwidth (OBW). This demonstrates a sharpening of tuning curves by inhibition, similar to the observation of Katzner et al. (2011), where inhibitory synapses in cat primary visual cortex were blocked with gabazine. Interestingly, PV cells seem not to affect the sharpening of tuning curves (Atallah et al., 2012; N. R. Wilson et al., 2012), whereas somatostatin-expressing neurons (SOM) sharpen neuronal responses (N. R. Wilson et al., 2012). This demonstrates the influences of the different inhibitory neuron types (Markram et al., 2004), which must be taken into account in future models.

### Shift in the E/I balance leads to the spontaneous emergence of contrast invariant tuning curves

As a consequence of the contrast gain mechanism by inhibition, our model shows the spontaneous emergence of contrast invariant orientation tuning (Skottun et al., 1987; Troyer et al., 1998; Finn et al., 2007). Early modeling studies have proposed feed-forward inhibition to implement a push-pull inhibitory mechanism for the emergence of contrast-invariant tuning curves (Troyer et al., 1998; Ferster & Miller, 2000). Despite the fact that our network contains feed-forward inhibition, we did not observe a push-pull inhibitory effect, in other words, anti-correlation of excitation and inhibition (Anderson et al., 2000). To be more specific, a direct comparison of the excitatory and inhibitory input current for the contrast invariance task shows a simultaneous increase and decrease of excitation and inhibition, caused by the detailed balance in our network (see supp. **Fig. S7**). We have observed that for the *EI*2/1 model, inhibitory input currents increase more rapidly than excitatory currents at higher contrast levels and non-preferred orientations. This results in a shift from a two-to-one ratio of excitation to inhibition to a one-to-one ratio between excitation and inhibition, and implements a contrast-dependent modulation of the neuron’s gain curve. In contrast to that, we observed for the *EI*3/1 model a proportional growth of the excitatory and inhibitory input currents for higher input contrast (see supp. **Fig. S8**), leading to an increase of the OBW. This shows that the emergence of contrast-invariant tuning curves is an inherent effect of the ratio between excitation and inhibition in our network, and suggests that contrast invariance emerge at a specific E/I ratio. A contrast-dependent shift in the balance between excitation and inhibition has been reported in the visual cortex of awake mice (Adesnik, 2017). Although the influence of inhibition on the neuronal gain function for the emergence of contrast invariance is in line with previous assumptions (Mitchell & Silver, 2003; Finn et al., 2007), recent studies have proposed that changes in the neuronal gain are caused by response variability in the afferent thalamic path (Sadagopan & Ferster, 2012; Priebe, 2016). An alternative proposal holds that fixed unspecific inhibition leads to contrast invariance (Ben-Yishai et al., 1995). We confirmed this by shuffling all synaptic weight to and from the inhibitory population. In this condition, we observed contrast-invariant tuning (see supp. **Fig. S10**). Our results extend these previous theories by showing that specific inhibition, as emerging through inhibitory plasticity and given sufficient inhibitory strength, is a sufficient condition for contrast invariance as well.

### Sparseness and metabolic efficiency benefit from E/I balance

We observed that in the *EI*2/1 model, the standard deviation of the membrane potential increases for non-preferred orientations. Together with the observed asynchronous spiking behavior, we conclude that the balance of inhibition and excitation leads to a more irregular spiking behavior. Previous work suggests that a more irregular activity and irregular membrane potential behavior is related to improved metabolic efficiency in terms of efficient input encoding (Deneve & Machens, 2016). Our observations agree with these findings, because the efficiency of information transmission in our network mainly benefits from the ratio between excitatory and inhibitory currents in the stable network.

An established approach in terms of input encoding efficiency is the concept of sparse coding (Rolls & Tovee, 1995; Vinje & Gallant, 2000; Tolhurst et al., 2009). However, in recent years, it has been discussed how the level of sparseness reported in physiological experiments is influenced by animal age and the level of anesthesia (Berkes et al., 2009), and the benefit of highly sparse codes for information processing has been questioned (Wiltschut & Hamker, 2009; Barak et al., 2013; Spanne & Jörntell, 2015). Overall, the intermediate sparseness values observed in our model are in agreement with experimental findings (Berkes et al., 2009; Froudarakis et al., 2014).

### Structured connectivity caused by inhibitory and excitatory plasticity

Previous physiological studies have shown that inhibitory interneurons are connected in a nonspecific manner to other cells in their surrounding (Harris & Mrsic-Flogel, 2013). However, recent studies observed that inhibitory PV cells develop strong connections to excitatory cells with similar orientations (Znamenskiy et al., 2018), and that neurons with similar preferred orientations have a higher probability for recurrent connections (Ko et al., 2011; Cossell et al., 2015).

We observed a similar connectivity pattern in our network, namely, the appearance of strong connectivity between co-tuned neurons. King et al. (2013) also obtained a structured connectivity between co-tuned excitatory and inhibitory cells in a spiking network. While King et al. (2013) achieved this goal by designing a suitable learning rule for the synaptic projections involving inhibitory neurons, we observed the appearance of strong connectivity as an emergent property of our model architecture based on detailed phenomenological rules.

### Stable learning despite limitations of simultaneous excitatory and inhibitory plasticity

Previous studies have mentioned the difficulty to achieve a certain level of inhibition in a network with inhibition and plastic excitatory synapses (Zenke & Gerstner, 2017; Hennequin et al., 2017). We next discuss the behavior of the selected learning rules more in detail to show some of the difficulties during the interaction of excitatory and inhibitory plasticity, and discuss the limitations of our modeling approach.

For the excitatory learning rule, Clopath et al. (2010) have shown that a lower input firing rate leads to bigger receptive fields, as a compensatory effect of the homeostatic mechanism. This mechanism is controlled by the long-term postsynaptic membrane potential in relation to a reference value and implements a local homeostatic mechanisms to influence the synaptic plasticity. If the membrane potential is too low, less long-term depression (LTD) in relation to long-term potentation (LTP) occurs, and the weights will increase. Otherwise, if the membrane potential is too high, a higher amount of LTD will occur to decrease the weights. Consequently, for a lower input firing rate, more weights will increase, saturating at their maximum, to achieve a specific postsynaptic activity.

The homeostatic mechanism of the inhibitory rule (Vogels et al., 2011) strengthens the inhibition if the postsynaptic activity is too high, with respect to a target firing rate (*ρ*), or decreases the weight otherwise. In our network, the postsynaptic membrane potential is a result of the difference between the incoming excitatory and inhibitory current, such that a reduction in the membrane potential through inhibition is comparable to a reduction through less presynaptic spikes. The operation of both homeostatic mechanisms on the postsynaptic activity leads to a competition between weight changes at excitatory and at inhibitory synapses and should lead to bigger receptive fields, or, in the worst case, to a saturation of all synapses to their maximum value (see supp. **Fig. S2** and **Fig. S3**).

However, we observed the emergence of stable receptive fields and stable connections between the populations. Additionally, our results show a reduction in the mean activity, caused by inhibition, without causing bigger receptive fields. We assume that in contrast to a reduction in the input, what leads to a proportional reduction on the postsynaptic neuron, the inhibitory current leads to a more irregular, or fluctuating, behavior of the membrane potential (Vogels et al., 2005). To allow LTP at excitatory synapses, the membrane potential must be higher than *θ*_+_ (= −45.3*mV*), which is slightly above the steady-state spiking threshold (*V_T_rest__* = −50.4*mV*). But if the membrane potential is hyperpolarized by inhibition, it falls below the LTP threshold: No LTP occurs, and the weights will not increase to the maximum. Additionally, we observed that the interplay of the excitatory and inhibitory rules are mainly influenced by the magnitude of learning rates. In particular, a higher excitatory or higher inhibitory learning rate led to the saturation of all synapses, as an effect of the competition between both homeostatic mechanisms. How fast the synaptic weight changes depends not only on the magnitude of learning rates, but also on the number of spikes, that is, the number of learning events. Therefore, the learning rates for the *noInh* model is smaller, to compensate the higher activity in the neuron populations. Finally, the competitive pressure between learning rules is controlled by the postsynaptic target activity in the inhibitory learning rule. Smaller values of *ρ* enhances the inhibitory pressure on the post-synaptic neuron to achieve a lower firing rate and can also lead to an unlimited growth of synaptic weights. This limited the amount of inhibition that can emerge in the network and did not allow a one-to-one balance between excitation and inhibition in our model, at least for natural scene stimuli. However, when presenting sinusoidal gratings of high contrast, E/I balance shifted towards a 1:1 ratio in the *EI*2/1 model, suggesting that this balance is stimulus-dependent.

Previous model studies reported that receptive fields can emerge without inhibition, by maintaining the post-synaptic activity over intrinsic plasticity (Savin et al., 2010), implementing a BCM-like behavior with a post-synaptic spike trace (Pfister & Gerstner, 2006), or regulating the LTD-term (Clopath et al., 2010). As expected by the chosen learning rules, our simulations with the *noInh* model confirm the emergence of receptive fields without inhibition. Despite this, other model studies pointed out the role of local homeostatic mechanisms on the emergence of selective receptive fields (Butko & Triesch, 2007; Zylberberg et al., 2011; Stevens et al., 2013) in networks with inhibition, or proposed that inhibition increases the diversity of receptive fields by implementing a competition between neurons (Földiak, 1990; Savin et al., 2010; King et al., 2013). In addition, our results show that plastic inhibition increases the receptive field diversity in comparison to fixed inhibition. By starting from unselective neurons, they develop a simple selectivity which pushes the correlation-based inhibitory influence to force a decorrelation between neurons and increase orientation divergence. This shows that inhibitory plasticity not only maintains the postsynaptic activity, but also implements a selective competition between neurons during a highly dynamical phase of development. Previous experimental studies mentioned different phases during the cortical maturation (Espinosa & Stryker, 2012; Toyoizumi et al., 2013; van Versendaal, Danielle and Levelt, Christiaan N., 2016), discussed the role of inhibition for the beginning of a critical period (Espinosa & Stryker, 2012; Toyoizumi et al., 2013), or showing a temporal decrease of inhibition to enable synaptic plasticity (van Versendaal, Daniölle and Levelt, Christiaan N., 2016). One of the best studied examples of critical period in the visual cortex is the onset of ocular dominance (OD) plasticity (Issa et al., 1999; Toyoizumi et al., 2013; van Versendaal, Daniölle and Levelt, Christiaan N., 2016). It has been discussed earlier that inhibitory interneurons (especially PV+) are important for the regulation of OD plasticity (Toyoizumi et al., 2013; van Versendaal, Danielle and Levelt, Christiaan N., 2016) and the strength of inhibition itself changes during this critical period (Gandhi et al., 2008; van Versendaal, Danielle and Levelt, Christiaan N., 2016), like a rapid downregulation of inhibitory cell activity (Kuhlman et al., 2013; van Versendaal, Daniölle and Levelt, Christiaan N., 2016). Our study about the role of inhibition for learning provides an excellent starting point for studies that aim to look at different critical periods in development.

### Conclusion

To the best of our knowledge, our simulations are the first demonstration of the parallel emergence of fundamental properties of the primary visual cortex such as sparse coding, contrast invariant tuning curves and high accuracy input representation, in a spiking network with spike timing-dependent plasticity rules. A central finding of our study is that the emergence of representational efficiency (such as tuning diversity) requires plasticity at feed-forward and feedback inhibitory synapses. Further, the emergence of a high tuning diversity as a direct consequence of inhibitory plasticity provides a verifiable prediction, via pharmacological or genetic methods, which allow to suppress inhibitory plasticity during the development of V1 simple cells. Although previous research has shown that unspecific inhibition has an effect on the gain-function of the excitatory cells, to improve the metabolic efficiency (Gaudry & Reinagel, 2007) or to cause contrast invariance (Ben-Yishai et al., 1995), our results demonstrate that the E/I ratio emerging from learning increases the metabolic efficiency (in terms of bits per spike) in our network. This emphasizes the role of inhibition in the shaping of neuronal responses (Isaacson & Scanziani, 2011; Stringer et al., 2016; Sprekeler, 2017) and in the development of reliable and efficient input encoding.

## Acknowledgment

This work has been supported by a grant from the European Social Fund (ESF) and the Free State of Saxony, and by a CRCNS US-German-Israeli collaboration on computational neuroscience grant “Multi-level neuro-computational models of basal ganglia dysfunction in Tourette syndrome” funded by the Federal Ministry of Education and Research, Germany (BMBF 01GQ1707), and in part by German Research Foundation (DFG, 416228727) - SFB 1410 Hybrid Societies.

## Author Contributions

Conceptualization F.H.H., M.T., L.G., R.L.

Methodology M.T., L.G., R.L.

Software R.L.

Validation R.L.

Formal Analysis R.L., L.G.

Investigation R.L., L.G., M.T.

Resources F.H.H.

Writing – Original Draft Preparation R.L., L.G.

Writing – Review & Editing F.H.H., L.G., M.T., R.L.

Visualization R.L.

Funding Acquisition F.H.H.

## 4 Methods

The first part of this section (4.1–4.5) describes the network architecture including the neuron model and learning rules. The model has been implemented in Python 3.6, using the ANNarchy simulator (Vitay et al., 2015), with a simulation time step of *dt* = 1*ms* (Euler integration). The neuronal simulator is available from https://bitbucket.org/annarchy/annarchy. The implementation of the adaptive exponential integrate-and-fire neuron model and the voltage-based triplet STDP learning rule from Clopath et al. (2010) based mainly on the re-implementation by Larisch (2019).

### 4.1 Network architecture

Our network model, which is inspired by the primary visual cortex and its inputs from LGN, consists of three populations of spiking neurons (**Fig. 1a**): An input layer representing LGN, and excitatory and inhibitory populations of V1, each receiving feed-forward inputs from LGN. The V1 populations are mutually interconnected via excitatory or inhibitory synapses, respectively. The circuit therefore implements both feed-forward and feedback inhibition, in agreement with anatomical findings (Isaacson & Scanziani, 2011). Inhibitory interneurons receive additional recurrent inhibitory connections. All projections follow an all-to-all connectivity pattern, excluding self inhibitory feedback connections.

The LGN layer consists of 288 neurons showing Poisson activity and is split into ON- and OFF-subpopulations. For the V1 excitatory population (144 neurons) and the inhibitory population (36 neurons), we used adaptive exponential integrate-and-fire neurons (Sec. 4.3). The size of the inhibitory population was chosen to match the 4:1 ratio between excitatory and inhibitory neurons found in visual and striate cortex (Beaulieu et al., 1992; Markram et al., 2004; Potjans & Diesmann, 2014). Researchers reported a much higher volume for the primary visual cortex than the LGN (Andrews et al., 1997), what suggests a much higher number of neurons. We verified tat the mere size of V1 in our model does not influence our conclusions, by increasing the number of excitatory and inhibitory cells by the factor of 2 and 10 using a sparse connectivity between excitatory and inhibitory cells to guaranty a similar E/I balance than for the *EI*2/1 model. We measured the input reconstruction error and the orientation bandwidth on different contrast levels and did not observe a high difference in comparison to the *EI*2/1 model (see **Fig. S12**).

All synaptic connections within our model are plastic and were randomly initialized. They change their weight based on either the voltage-based STDP-rule proposed by Clopath et al. (2010) (excitatory connections) or the symmetric iSTDP-rule proposed by Vogels et al. (2011) (inhibitory connections; Sec. 4.5).

Although networks of the visual cortex have lateral excitatory connections (Ko et al., 2011, 2013; Harris & Mrsic-Flogel, 2013; Lee et al., 2016) as also discussed in different model studies (Litwin-Kumar & Doiron, 2014; Miconi et al., 2016; Sadeh et al., 2015), we did not insert plastic lateral excitatory connections in our model, as our model is already highly adaptive and further excitatory connections may complicate the required set of learning rules. However, to observe the influence of lateral excitation, we inserted fixed excitatory connections with a connection probability of 0.2 between the excitatory neurons and initialized the weight with an uniform distribution. Despite the number of unstable learning approaches increased (see **Fig. S13**), we did not observe a significant influence of the recurrent connections, by measuring the IRE and OBW.

### 4.2 Network input

As network input, we used whitened patches from natural scenes (Olshausen & Field, 1996, 1997). Each patch was chosen randomly, with a size of 12 by 12 by 2 pixels (Wiltschut & Hamker, 2009). The third dimension corresponds to the responses of ON- and OFF-cells. To avoid negative firing rates, we mapped positive pixel values to the ON-plane, and the absolute value of negative pixels to the OFF-plane. Every patch was normalized with the maximum absolute value of the corresponding natural scene. The firing rate of each Poisson neuron represents the brightness value of the input pixels. The firing rate associated to the (rarely occurring) maximum pixel value was set to 125*Hz*. We stimulated the network with 400.000 patches during training, with a presentation time of 125*ms* per patch, corresponding to around 14*h* of simulated time. To avoid any orientation bias in the input, the patch was flipped around the vertical or horizontal axis independently with 50% probability (Clopath et al., 2010).

### 4.3 Poisson neuron model in LGN

For modeling convenience, we generated Poisson activity in LGN neurons by injecting brief voltage pulses, generated by a Poisson process, into a binary spiking neuron model, such that each voltage pulse input triggered a spike. This simplified the numerical calculation of a spike trace required for the learning rule, while preserving the precise timing of spikes drawn from a Poisson process.

The spike trace 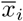 is updated whenever the presynaptic neuron *i* spikes, and decays exponentially: *X_i_*(*t*) = 1 if a spike is present at time *t*, and *X_i_*(*t*) = 0 otherwise.

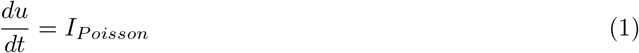

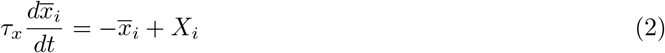

### 4.4 Adaptive exponential integrate-and-fire neurons in V1

For the neurons in the V1 excitatory and inhibitory layer, we used a variant of the adaptive exponential integrate-and-fire model as described by Clopath et al. (2010). In this model, the membrane potential *u* is influenced by the following additional dynamical variables: An adaptive spike threshold, *V_T_*, a hyperpolarizing adaptation current, *w_ad_*, and a depolarizing afterpotential, *z*. Excitatory and inhibitory synaptic currents are denoted by *I_exc_* and *I_inh_*. For an explanation of constant parameter values as used by Clopath et al. (2010), see Table 1.

The full equation for the membrane potential is

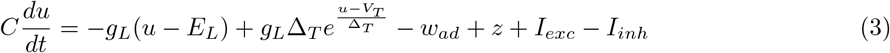

**Table 1:**
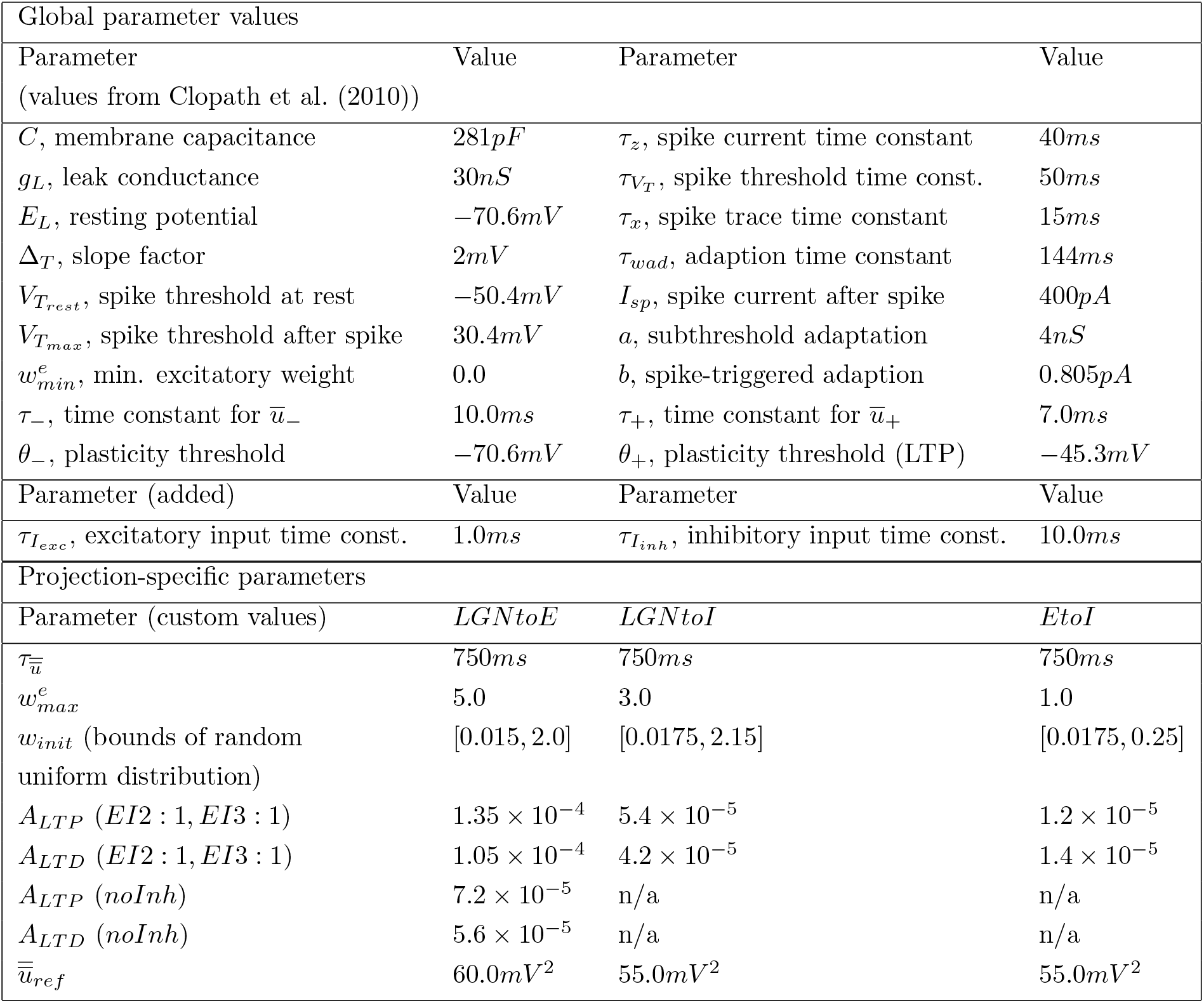
Parameters for the neuron model and excitatory synapses. Note that for the *noInh* model, learning rates were reduced to compensate for the increased firing rates in the absence of inhibition.

As the triplet voltage STDP rule is sensitive to the precise time course of the membrane voltage, including the upswing during a spike, the magnitude of weight changes depends on the implementation details of the after-spike reset. To avoid long simulation times associated with smaller time steps, we opted for the following simplified treatment of the spike waveform which reproduced the results reported by Clopath et al. (2010): Whenever the membrane potential *u* exceeded the spike threshold, *u* was held at a constant value of 29*mV* for 2*ms*, and then reset to the resting potential *E_L_*. We obtained highly similar results from an alternative implementation, in which the after-spike reset was immediately applied when the spike threshold was crossed, with an additional update of the voltage traces by the amount expected from a 2*ms*-long spike (data not shown).

The reset value for the spike threshold is *V_T_max__*, with exponential decay towards the resting value *V_T_rest__*, with a time constant *τ_V_T__* (Eq. 4):

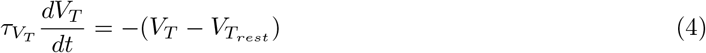

The afterpotential *z* has a reset value of *I_sp_* and decays to zero (Eq. 5). Further, the variable *w_ad_* is incremented by the value *b* and decays exponentially (Eq. 6).

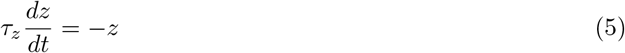

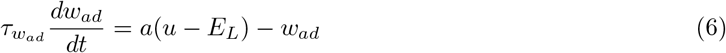

The model proposed by Clopath et al. (2010) assumed excitatory synaptic input in the form of voltage pulses. For modeling convenience, we approximated this setting by current-based excitatory synapses with a short time constant of 1*ms*. Inhibitory synaptic currents decayed with a slower time constant of 10*ms*. Both synaptic currents are incremented by the sum of synaptic weights of those presynaptic neurons which spiked in the previous time step:

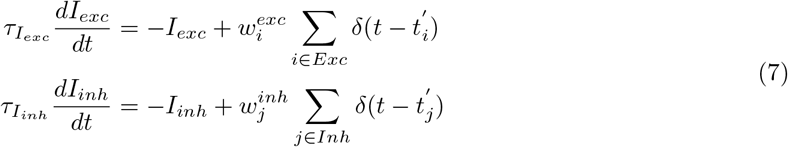

where 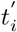 denotes the spike time of presynaptic neuron *i*, and *δ* is the indicator function with *δ*(0) = 1.

### 4.5 Synaptic plasticity

#### 4.5.1 Voltage-based triplet STDP at excitatory synapses

Plasticity at excitatory connections (LGN to Exc. and Exc. to Inh.) follows the voltage-based triplet STDP rule proposed by Clopath et al. (2010). We here repeat the essential features of this plasticity model. The neuronal and synaptic variables describing the development of the weight from a presynaptic neuron with index *i* onto a given postsynaptic neuron are: *X_i_*, the presence of a presynaptic spike; 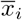, the presynaptic spike trace (Eq. 2); *u*, the postsynaptic neuron’s membrane potential; and two running averages of the membrane potential, 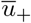 and 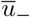, defined as follows:

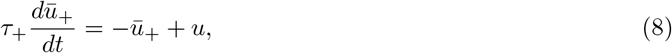

where 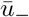 is defined analogously, with the time constant *τ_−_*. In addition, the learning rule includes a homoeostatic term, 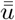, which regulates the relative strength of LTD versus LTP, and which measures the mean postsynaptic depolarization on a slower time scale:

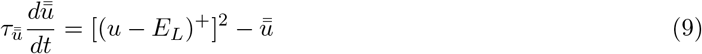

Here, *x*^+^ = max(*x*, 0) denotes top-half rectification.

The full learning rule is given as the sum of the LTP term and the LTD term:

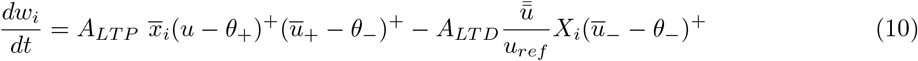

where *A_LTP_* and *A_LTD_* are the learning rates for LTP and LTD, *θ*_+_ and *θ*_−_ are threshold parameters, and *u_ref_* is a homeostatic parameter which controls the postsynaptic target firing rate. Clopath et al. (2010) have shown that this learning rule results in BCM-like learning dynamics (Bienenstock et al., 1982), in which a sliding metaplasticity threshold leads to the development of selectivity.

Following Clopath et al. (2010), for the LGN efferent connections, we equalized the norm of the OFF weights to the norm of the ON weights every 20*s*. The weight development is limited by the hard bounds 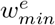 and 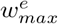.

**Table 2:** Parameters for inhibitory synapses.

|  | ItoE and ItoI | ItoE | ItoI |
| --- | --- | --- | --- |
| $\tau_{post}$ | 10.0ms | | |
| $\tau_{pre}$ | 10.0ms | | |
| $w^i$ initial | 0.0 | | |
| $w_{min}^i$ | 0.0 | | |
| $w_{max}^i$ | | 0.7 | 0.5 |
| $\eta$ | | $10^{-5}$ | $10^{-5}$ |
| $\rho$ (EI3 : 1) | | 0.7 | 0.6 |
| $\rho$ (EI2 : 1) | | 0.4 | 0.6 |

#### 4.5.2 Homeostatic inhibitory plasticity

Previous biological studies have observed spike timing-dependent plasticity of inhibitory synapses which differs from the well-known asymmetric STDP window (Caporale & Dan, 2008; D’Amour & Froemke, 2015). We therefore chose to implement the phenomenologically motivated, symmetric inhibitory STDP (iSTDP) rule proposed by Vogels et al. (2011) at all inhibitory synapses (Eq. 11):

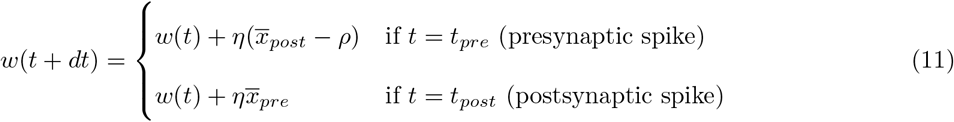

Here, *η* is the learning rate, and *ρ* is a constant which controls the amount of LTD relative to LTP. Further, Vogels et al. (2011) have shown that this learning rule has a homeostatic effect, and the parameter *ρ* controls the postsynaptic target firing rate. The variables 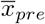 and 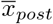 are spike traces for the pre- and postsynaptic neurons, defined in analogy to Eq. (2), with time constants *τ_pre_* and *τ_post_*. In this plasticity rule, near-coincident pre- and post-synaptic spiking causes potentiation of weights, irrespective of their temporal order. By contrast, isolated pre- or postsynaptic spikes cause depression of weights. As for the excitatory learning rule, weights are bounded by 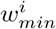 and 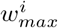. For parameter values, see Table 1.

#### 4.5.3 Choice of parameter configurations

As our main goal is to determine the influence of inhibitory strength both on the formation of selectivity and on the dynamics of stimulus coding, we simulated our network using different parameter and network configurations. First, we used the above presented network, where the strength of the inhibitory feedback is controlled by the homeostatic parameter *ρ*. With *ρ* = 0.4 for the feedback inhibitory synapses, we achieved a ratio of excitation to inhibition (E/I-ratio) of approximately 2: 1 on patches of natural scenes (abbreviated as *EI*2/1). On one hand, a lower *ρ* would strengthen the inhibitory feedback, but caused unstable behavior during learning. On the other hand, a higher *ρ* would weaken the inhibitory feedback of the model. Because of this, we were unable to achieve a 1: 1 E/I ratio for natural scene patches. With *ρ* = 0.7 we achieve a E/I-ratio of approximately 3: 1 on natural scene input (abbreviated as *EI*3/1), this led to similar but weaker characteristics for most of the experiments (**Fig. 1b**).

Second, we simulated a purely excitatory feed-forward network without any inhibitory activity (abbreviated as *noInh*), as the learning rule proposed by Clopath et al. (2010) is capable of learning distinct shapes of receptive fields given different initial weights.

Further, to control for the dynamical effects of inhibition in the steady state following receptive field development, we simulated the effects of deactivating the inhibitory synaptic transmission in the *EI*2/1 model after learning (abbreviated as *blockInh*). All three model variations are based on the same network architecture, consisting of the same number of neurons in each population and the same number of synapses, except that inhibitory weights differ in their strength or are deactivated. The different parameters for learning the models are shown in Table 1. We took the parameters for the adaptive integrate and fire neuron from Clopath et al. (2010). Based on the original parameter mentioned in Clopath et al. (2010) and Vogels et al. (2011), the parameters for both learning rules were found empirically to enable a stable emergence of receptive fields in multiple runs, initialized with different weight values (see supp. **Fig. S11**).

To test the stability and the reproducibility of our results, we performed 20 runs of each model with randomly initialized synaptic weights.

To evaluate how inhibitory plasticity interacts with plastic excitation, we deactivated the plasticity for specific synapses for three model variations. First, we deactivated the plasticity only in the inhibitory feedback connections (*fix fb inh*). Second, the plasticity was deactivated in both excitatory connections the inhibitory population (*fix ff inh*). We further deactivated the plasticity in the connections from the excitatory to the inhibitory population and for the lateral inhibition. Additionally, we trained one model variation where all connections were plastic, to validate that the learning is successful with pre-trained, shuffled weight matrices. To ensure that the same average amount of excitatory or inhibitory current is conveyed by the fixed synapses, we used shuffled weight matrices from previous simulations of the *EI*2/1 model for the respective synapses. No parameter changes were needed. To test the stability and reproducibility, we performed five runs of each variation.

### 4.6 Analysis methods

#### 4.6.1 Receptive field mapping

Over the course of learning, the excitatory input weights from LGN to V1 develop based on the pre- and postsynaptic activity. It is therefore possible to obtain a good approximation of the neurons’ receptive fields (RFs) by taking the weight matrix and reverting the ON-OFF mapping. To do this, we subtract the OFF-synapses from the ON-synapses and receive the receptive field. This is possible as either the ON- or the OFF-synapses can be activated by the input, so that the weights will also follow this distribution.

In addition to the visualization based on weight matrices, the receptive fields can also be revealed by probing the neurons with random stimuli. This approach has been successfully used in physiological research, in form of the spike triggered average (STA) (Ringach & Shapley, 2004; Schwartz et al., 2006; Pillow & Simoncelli, 2006). In this method, a neuron’s receptive field is defined as the average of white noise stimuli, weighted by the stimulus-triggered neuronal activity. We applied this method on the learned neural network. We presented noise patches drawn from a normal distribution with *μ* = 15, *σ* = 20 as input image to the network, and converted these to Poisson spike trains (cf. Sec. 4.2). Negative pixel values were set to zero, and the presentation time per patch was 125*ms*. For each neuron, we recorded the number of spikes per stimulus and calculated the average across all stimuli, weighted by the number of postsynaptic spikes (Eq. 12)

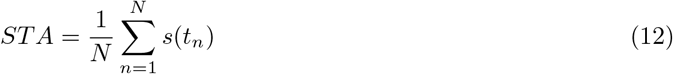

Here, *s*(*t_n_*) is the input stimulus at time point *t_n_*, when the nth spike has occurred, and *N* is the total number of postsynaptic spikes. Accordingly, stimuli evoking more spikes are higher weighted than stimuli evoking few or no spikes.

As we observed a high similarity between each neuron’s STA and its ON-OFF receptive field, we concluded that the overall receptive field shape was not significantly influenced by inhibition. Thus, for simplicity, the feed-forward weight vectors can be used for further evaluations.

#### 4.6.2 Receptive field similarity

As mentioned above, the feed-forward weight vector approximates the receptive field of a neuron. To measure the similarity between two receptive fields, we calculate the cosine between their feed-forward weight vectors (Eq. 13).

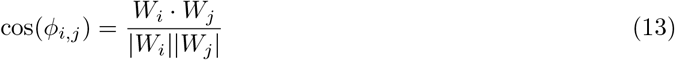

A value near +1 indicates high similarity, values around zero describe orthogonal weight vectors, and values near −1 indicates inverted weight vectors (i.e., maximally overlapping RFs with opposite directional preference).

#### 4.6.3 Tuning curves and orientation selectivity

The orientation selectivity is a well-studied characteristic of simple cells in V1 of mammals (Gilbert & Wiesel, 1990; Priebe & Ferster, 2008; Niell & Stryker, 2008) and thus, also a topic of interest for models of the visual cortex (e.g., Sadeh et al., 2014; W. Zhu et al., 2010; Tao et al., 2004). One possibility to quantify the orientation selectivity of a neuron is to measure its tuning curve (Ringach et al., 2002). For simple cells in the primary visual cortex, the orientation tuning curve describes the magnitude of responses evoked by a stimulus presented at different angles. In many biological studies, the tuning curves have been measured based on two-dimensional sinusoidal gratings (Anderson et al., 2000; Smith & Kohn, 2008; Ringach et al., 2002; Katzner et al., 2011). Therefore, we measured the responses to sinusoidal grating stimuli, rotated in steps of 8°, with different spatial phases from *0rad* to *πrad*, a different spatial frequencies from 0.05 up to 0.15*cycles/pixel*, centred to the input space and with a presentation time of 125*ms*.

Because of Poisson activity in the input layer, neuronal activity shows trial-to-trial fluctuations. Hence, we repeated every presentation 50 times, and calculated the mean across all 50 repetitions (or 6.25*s* presentation time). In contrast to the natural scene input used for training, the maximum input firing rate was set to 85.7*Hz*. This was suitable to obtain sufficiently high activity levels.

To estimate tuning curve sharpness, we calculated the orientation bandwidth (OBW) for every neuron. The OBW is defined as the half-width of the tuning curve, at an activity level of 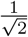 (approx. 70.7%) of the maximum (Ringach et al., 2002). Higher OBW values correspond to a broader tuning curve, and vice versa. Other definitions use the height at half-maximum, which does not change the overall result of this evaluation.

#### 4.6.4 Orientation diversity

To quantify the diversity of receptive field orientations, we calculated a histogram over the measured preferred orientations to measure the distribution and the incidence of a specific orientation (*P*(*o*) where *o* is the index to a specific orientation) Then, we calculated the Kullback-Leibler divergence (Eq. 14) between this distribution and an idealized uniform distribution of orientations (*Q*(*o*)).

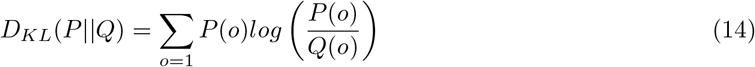

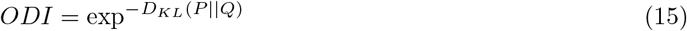

To calculate the orientation diversity index (*ODI*), we used the exponential function on the calculated Kullback-Leibler divergence. A value closer to one indicates a more uniform distribution of the measured orientations and thus a higher orientation diversity, whereas a value closer to zero indicates a less uniform distribution and thus a lower orientation diversity.

#### 4.6.5 Neuronal gain curves

A neuron’s gain function describes how neuronal activity is scaled by variations in the magnitude of excitatory inputs (Katzner et al., 2011; Isaacson & Scanziani, 2011). While an integrate-and-fire neuron receiving only excitatory inputs has a relatively static gain function (also called transfer function), controlled by the parameters of the neuron model, additional inhibitory inputs can modulate the effective input-to-output relationship. To characterize these inhibitory influences on gain curves, we recorded the excitatory synaptic currents and spiking activity evoked by sinus gratings (see Sec. 4.6.3), which we rotated from the orthogonal towards the preferred orientation of each neuron. Further, we changed the contrast of the input, by changing the pixels relative to the maximum input firing from 14.25*Hz* up to 100*Hz*. As before, we presented each stimulus orientation for 125*ms*, repeated 50 times (6.25s), and determined gain curves based on the average spike count across these 50 repetitions. We measured the spike count for each input degree and contrast strength and sorted the neuronal activity to the corresponding excitatory input, in ascending order.

#### 4.6.6 Measurement of E to I ratio

To determine the ratio between excitatory and inhibitory input current, we measure both incoming currents for the excitatory population for 1.000 randomly chosen natural scenes. Every scene was presented for 125*ms* and was repeatedly shown for 100 times. We averaged the incoming currents over the input stimuli repetitions and sorted for each neuron and stimuli the excitatory input currents ascending with the related inhibitory currents. For better visualization, the currents are summarized into bins.

#### 4.6.7 Sparseness

The sparseness value expresses the specificity of population codes and single neurons, both in experimental studies (Rolls & Tovee, 1995; Vinje & Gallant, 2000, 2002; Weliky et al., 2003; Tolhurst et al., 2009) and in model simulations (Wiltschut & Hamker, 2009; Zylberberg et al., 2011; King et al., 2013). It quantifies either the fraction of neurons which respond to a single stimulus, called population sparseness, or the number of stimuli to which a single neuron responds, called lifetime sparseness (Tolhurst et al., 2009). In the past, many different sparseness measurements are established (Rolls & Tovee, 1995; Hoyer, 2004). To measure the specificity of our network activity, we calculated the population sparseness after Vinje & Gallant (2000) (see Eq. 16).

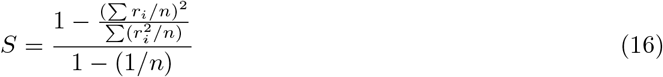

where *r_i_* is the activity of the *i*th neuron to a specific input and *n* the number of neurons in the neuron population.

By construction, sparseness values are bound between zero and one. If the neuron population has dense activity, i.e., most neurons are active to an input stimulus, the sparseness level approaches zero. By contrast, few active neurons of the population lead to a sparseness value close to one. As input, we used 30.000 natural scene patches, and determined sparseness values based on the firing rates of each neuron on each input patch.

#### 4.6.8 Image reconstruction error

The network’s coding performance following training can be measured by the difference between input images and their reconstruction from network activity. This method gives direct insight on how well visual input is represented by the network as a whole. This aspect was often not considered in previous biologically motivated circuit models of the primary visual cortex. We used the root mean-square error between one image of the natural scenes dataset from Olshausen & Field (1996) and the reconstructed one (cf. Zylberberg & DeWeese, 2013; King et al., 2013) (Eq. 17), termed image reconstruction error (IRE):

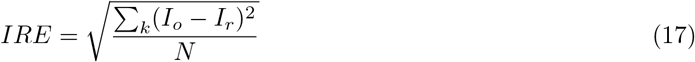

where *N* denotes the number of image pixels. To obtain the reconstructed image *I_r_*, we subdivided the full image into patches of size 12 × 12, in an overlapping fashion (in increments of 3 pixels). We showed each patch 50 times for 125*ms* each, and recorded neuronal activities. We weighted the activity of each neuron by its feed-forward weights to obtain a linear reconstruction of each image patch, which we combined to reconstruct the full image. This approach is equivalent to calculating the IRE for individual patches, and calculating the root mean-square of these individual IRE values. To ensure that pixel values of the reconstructed image were in the same range as the original image, we normalized the reconstructed as well as the original image to zero mean and unit variance (Zylberberg & DeWeese, 2013; King et al., 2013).

#### 4.6.9 Mutual information

We measure the metabolic efficiency via the numbers of spikes which are necessary to represent a specific input stimuli and the amount of information transmitted via a spike. An information-theoretic approach to estimate this coding efficiency of the network is based on the mutual information between stimulus identity and neuronal activity (Dayan & Abbott, 2001; Dadarlat & Stryker, 2017). This measure allows to calculate the average information transmission per spike (Vinje & Gallant, 2002; Sengupta et al., 2013). To quantify information transmission, we calculated the mutual information, *I*(*s, r*), between the stimulus identity and neuronal responses for each neuron, following Vinje & Gallant (2002):

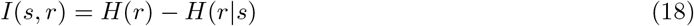

In Eq. 18, *I*(*s, r*) is the mutual information carried between stimulus and response for a time bin of 125*ms* length, the duration of a single stimulus. For that purpose, we calculate the total response entropy, *H*(*r*), and the conditional response entropy, also called stimulus-specific noise entropy, *H*(*r|s*).

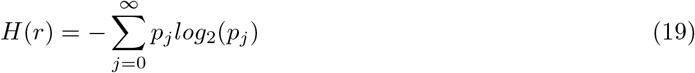

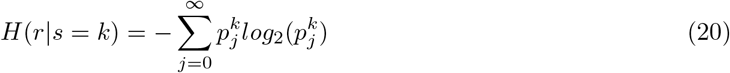

The total response entropy is given by Eq. 19. The variable *p_j_* is the number of time bins containing exactly *j* spikes, divided by the total number of time bins, or stimuli. It follows from Eq. 19 that the total response entropy is maximal if all spike counts occur with equal probability (and, if they do, the number of possible spike counts increases the entropy). The noise entropy for a specific stimulus (see Eq. 20) describes the variability of the neuronal responses across repetitions of a single stimulus *k*. Every stimulus was repeated 100 times. Similar to the total response entropy, *j* is the number of spikes which occurred in response to a stimulus *k*. Here, 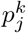 is the number of repetitions of stimulus *k* to which exactly *j* spikes are emitted, divided by the overall number of repetitions of that stimulus. To calculate the overall noise entropy of a neuron *H*(*r|s*), we averaged the noise entropy across all stimuli. Information per spike was computed by dividing *I*(*s,r*) by the mean number of spikes per stimuli, or time bins.

#### 4.6.10 Discriminability

To evaluate how well the network responses allow to distinguish between any two input patches, in the presence of trial-to-trial (how much is the variance in the firing rate of a neuron to specific input (Shadlen & Newsome, 1998)) fluctuations induced by Poisson input, we calculated the discriminability index, *d′* (e.g., Dayan & Abbott, 2001; Dadarlat & Stryker, 2017). The *d′* index measures the separation of two random distributions, and is closely related to the performance of a linear classifier assuming independent neuronal responses. Based on a random set of 500 natural scene patches, we calculated the *d′* by pairing the response on every patch to all other patches. For each pair of stimuli, *s*_1_ and *s*_2_, we presented each stimulus with *N* = 100 repetitions, and recorded the network responses of all *n* = 144 excitatory neurons for each repetition, obtaining the n-dimensional response vectors 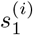 and 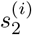, *i* = 1,…, *N*. We first calculated the mean activity of each cell in response to each stimulus, across the *N* repetitions (denoted by 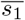 and 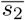). We next projected each individual population response 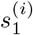 and 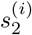 onto the vector between these means, by taking the dot product between each response and the difference 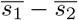:

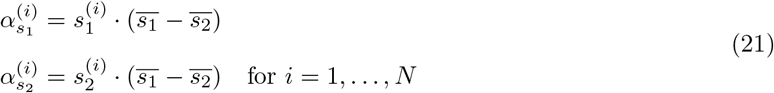

where *α_s_1__* and *α_s_2__* denote the projected responses. Next, we calculated the means and variances of the projected responses *α_s_1__* and *α_s_2__*, denoted by 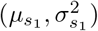 and 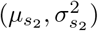. Finally, we calculate the discriminability 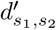, as the ratio between the separation of the means and the variances of the projected data:

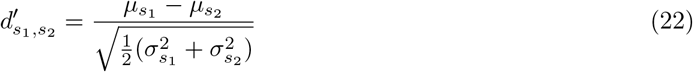

Note that we used the same sequence of patches for all model configurations to calculate the discriminability, and every patch was presented for 125*ms*. Previous research found that the variance of the response of a neuron to input stimuli is proportional to the mean (Gershon et al., 1998). Further studies demonstrated that inhibition leads to less variance in the responses to one repeatedly shown stimulus (Haider et al., 2010). The discriminability (*d′*) increases if the response variance decreases by the same response mean. Therefore, we can measure differences in the response variance.

