## Supplementary for "Sensory coding and contrast invariance emerge from the control of plastic inhibition over emergent selectivity"

**Supplementary Material**

René Larisch [1,*], Lorenz Gönner [1,2], Michael Teichmann [1] and Fred H. Hamker [1,3,**]

August 23, 2021

[1] *TU Chemnitz, Dept. of Computer Science, Artificial Intelligence*
[2] *TU Dresden, Faculty of Psychology, Lifespan Developmental Neuroscience*
[3] *Bernstein Center Computational Neuroscience Berlin*
[*] **
[**] **

### 1 Supplementary material

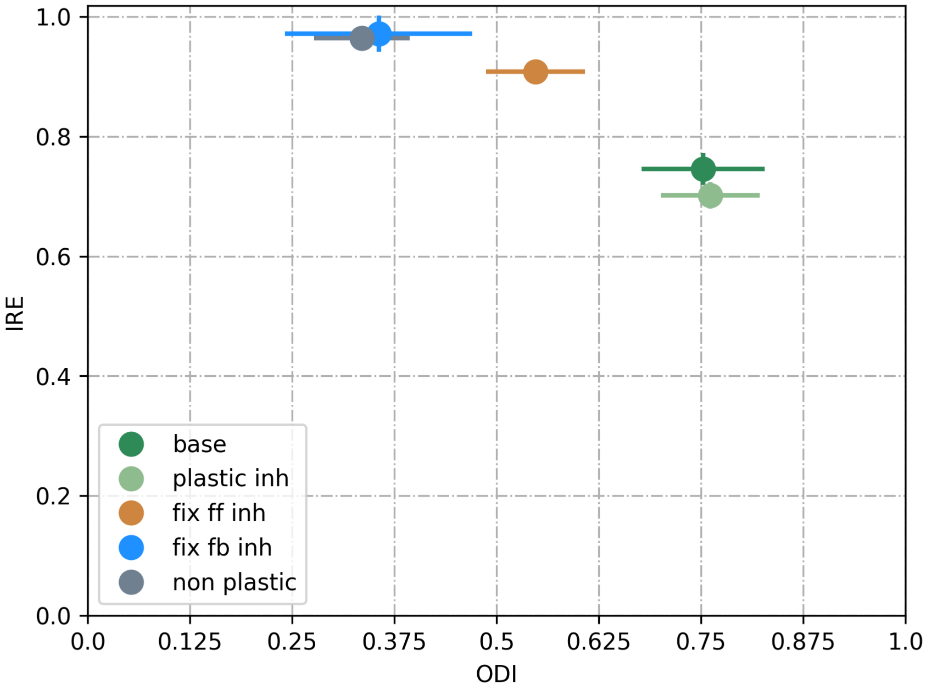


S 1: Image reconstruction error (IRE) for different fixed and plastic inhibitory connections. Excitatory synapses learned with the Clopath et al. (2010) learning rule. The dark green model (called *base*) is equal with the $EI2/1$ model. The other models are initialized with shuffled weights of a previous successfully learned $EI2/1$ model. In the *plastic inh* model, all inhibitory synapses are plastic, in the *fix ff inh* model is the feed-forward inhibition fixed, in the *fix fb inh* model is the feedback inhibition fixed, and in the in the *non plastic* model are all inhibitory connections fixed.

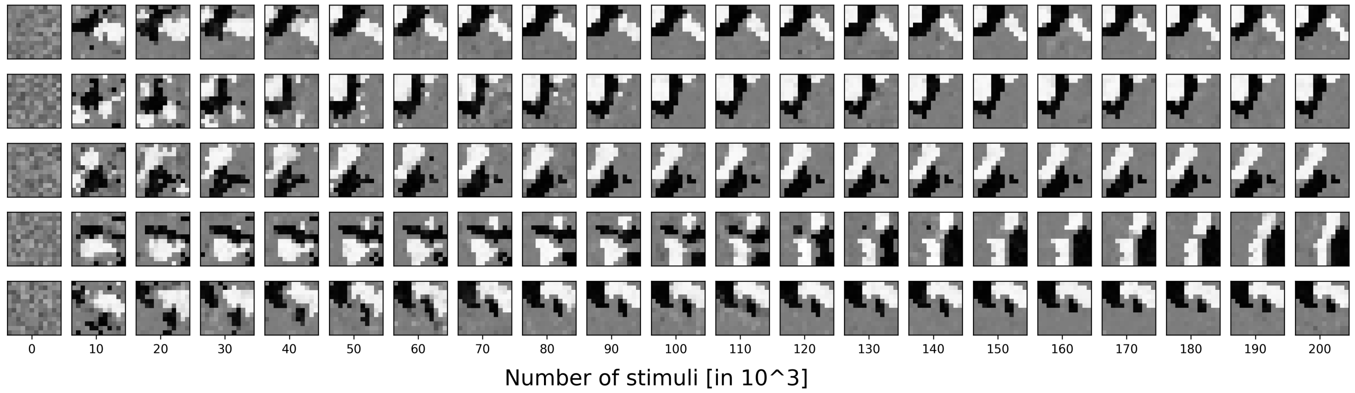


(a)

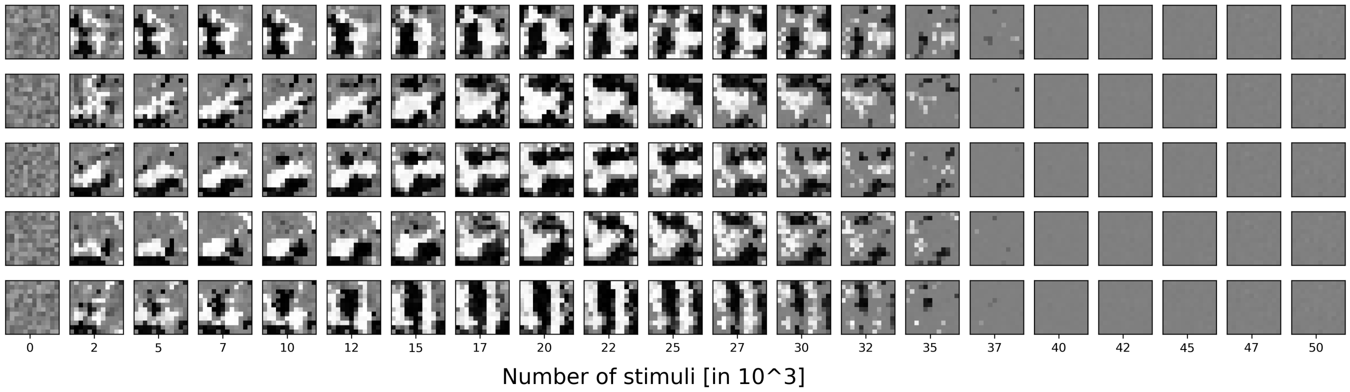


(b)

S 2: **Development of receptive fields.** Input weights of five randomly chosen excitatory cells. Bright values show input from the ON-LGN population and dark values from the OFF-LGN population. ON and the OFF weights are subtracted from each other to show the receptive fields. **(a)** Emergence of stable receptive fields. **(b)** Examples of unstable receptive fields, i.e when the differences between ON and OFF weights are zero (gray values) due to the increase of both components to the maximum value.

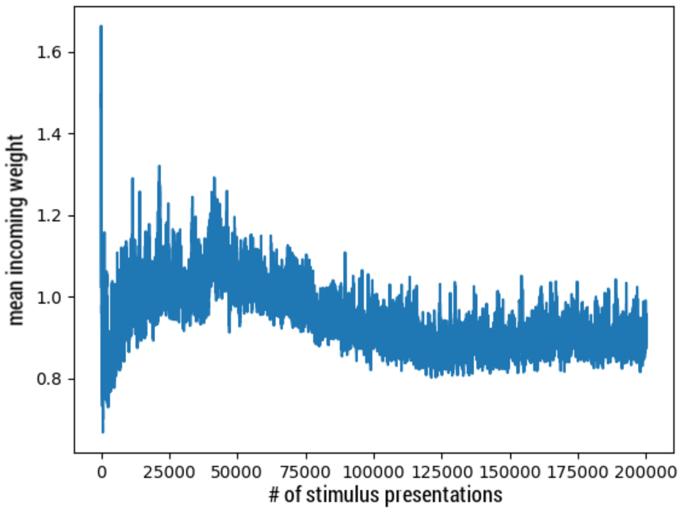


(a)

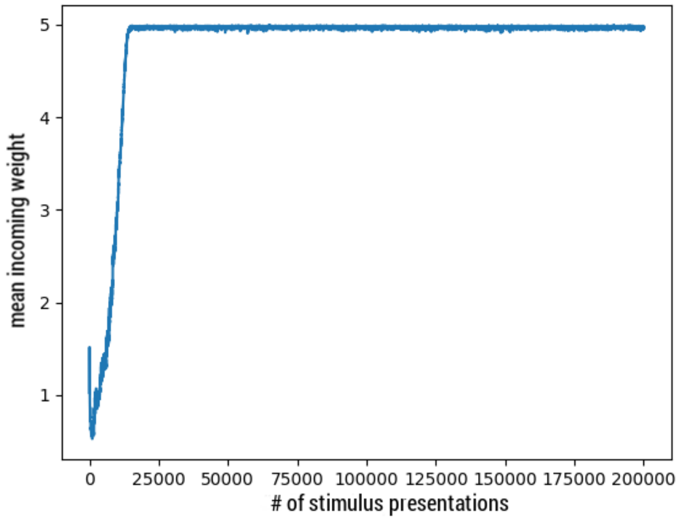


(b)

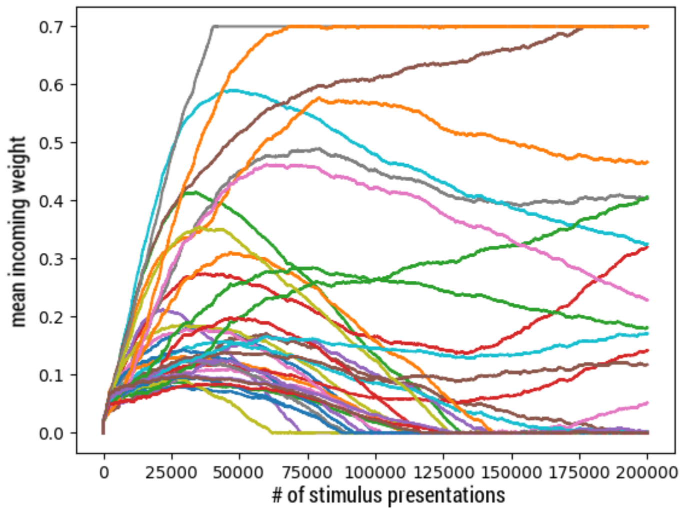


(c)

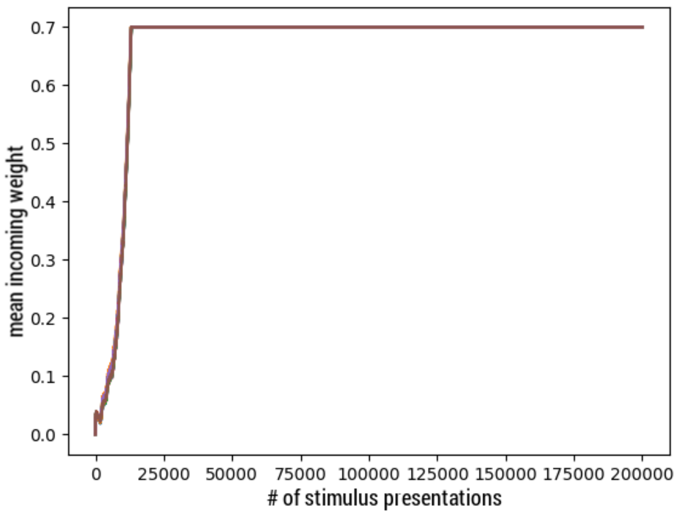


(d)

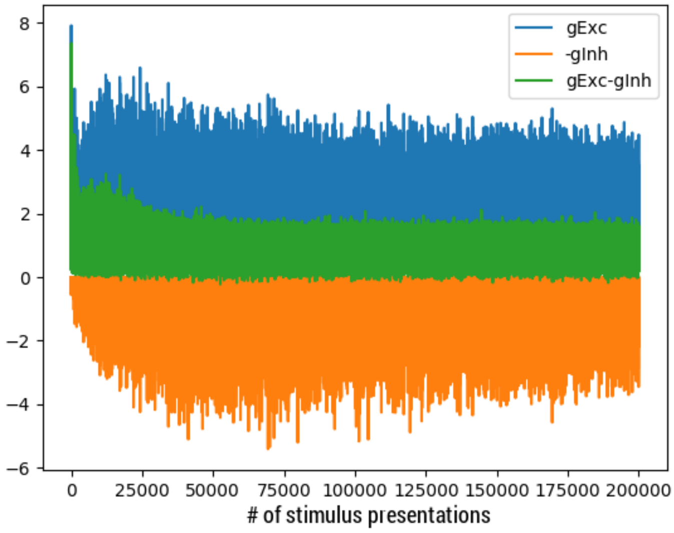


(e)

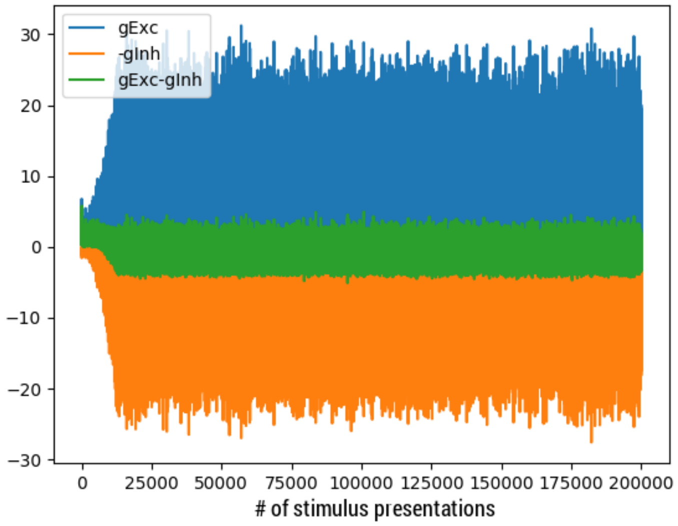


(f)

S 3: **Dynamics of weights during the first** $200,000$ **stimulus presentations.** First column shows stable weight learning. Second column shows the unlimited growth of the weights against the maximum weight value. **(a)** and **(b)** mean feed-forward excitatory weights from the LGN population to one excitatory neuron. **(c)** and **(d)** inhibitory feedback weights from the inhibitory population to one excitatory neuron. **(e)** and **(f)** mean input currents of the excitatory population.

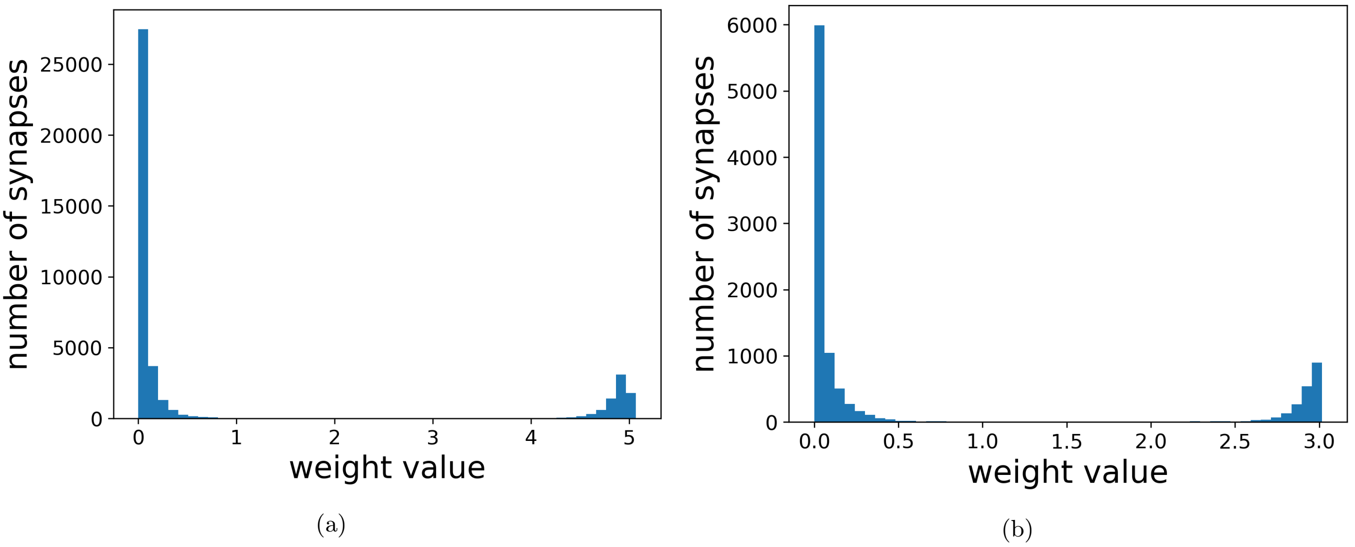


S 4: Histogram of feed-forwards weights from the LGN to the excitatory population **(a)**, and of the feed-forwards weights from the LGN to the inhibitory population **(b)** from the $EI2/1$ model.

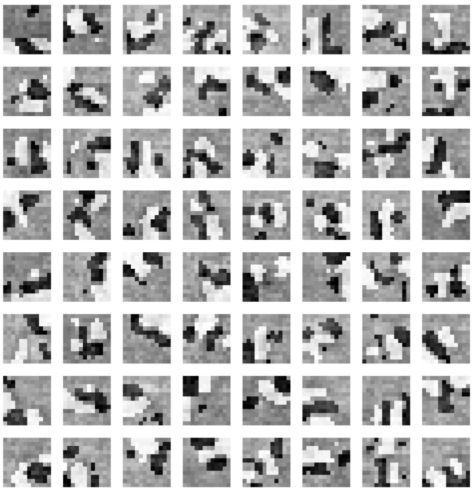

(a)

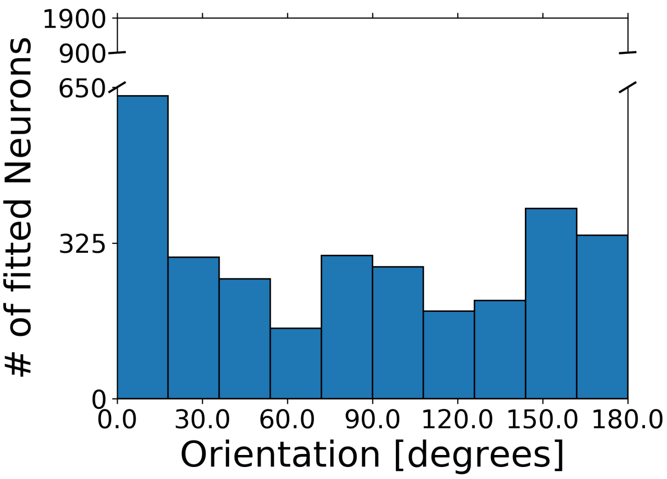


(b)

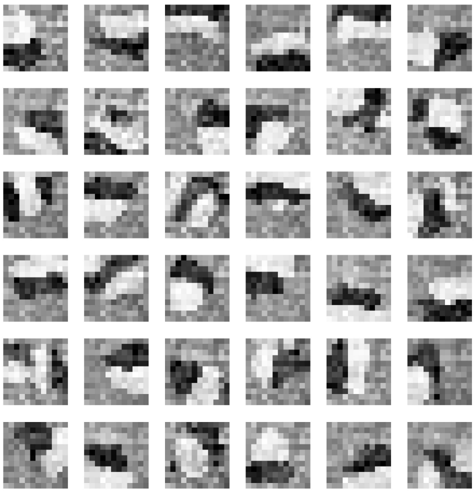

(c)

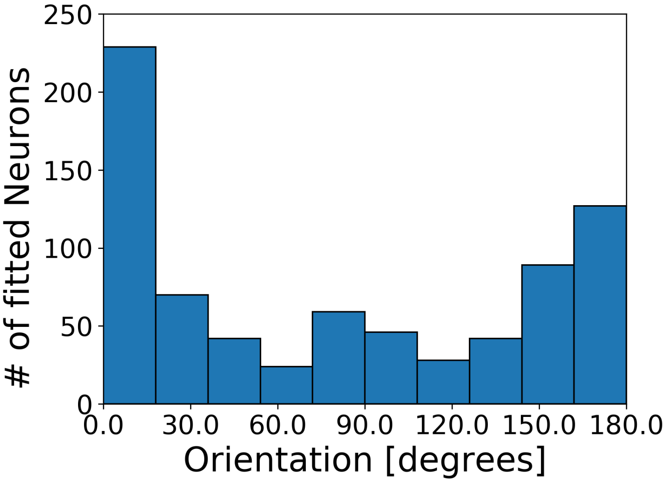


(d)

S 5: **(a)** Receptive fields of randomly selected 64 excitatory neurons of the $EI3/1$ model. **(b)** Distribution of receptive field orientation of all excitatory neurons of 20 model runs ($EI3/1$ model). **(c)** Receptive fields of all 36 inhibitory neurons of the $EI3/1$ model. **(d)**Distribution of receptive field orientation of all inhibitory neurons of 20 model runs ($EI3/1$ model) ($EI3/1$ model)

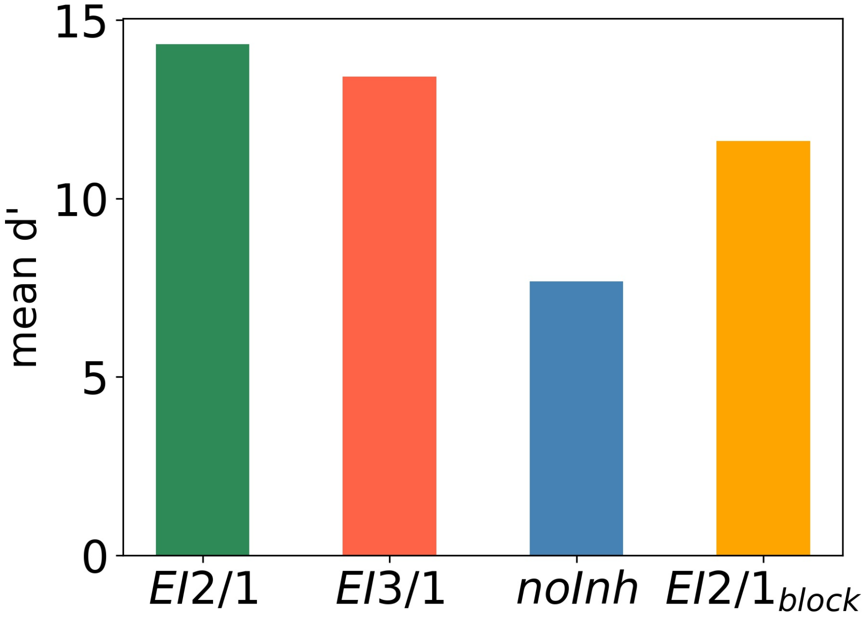


S 6: Average discriminability ($d'$) based on the responses to 500 randomly chosen natural scene patches. Discriminability benefits from tuning diversity of receptive fields and from feedback inhibition.

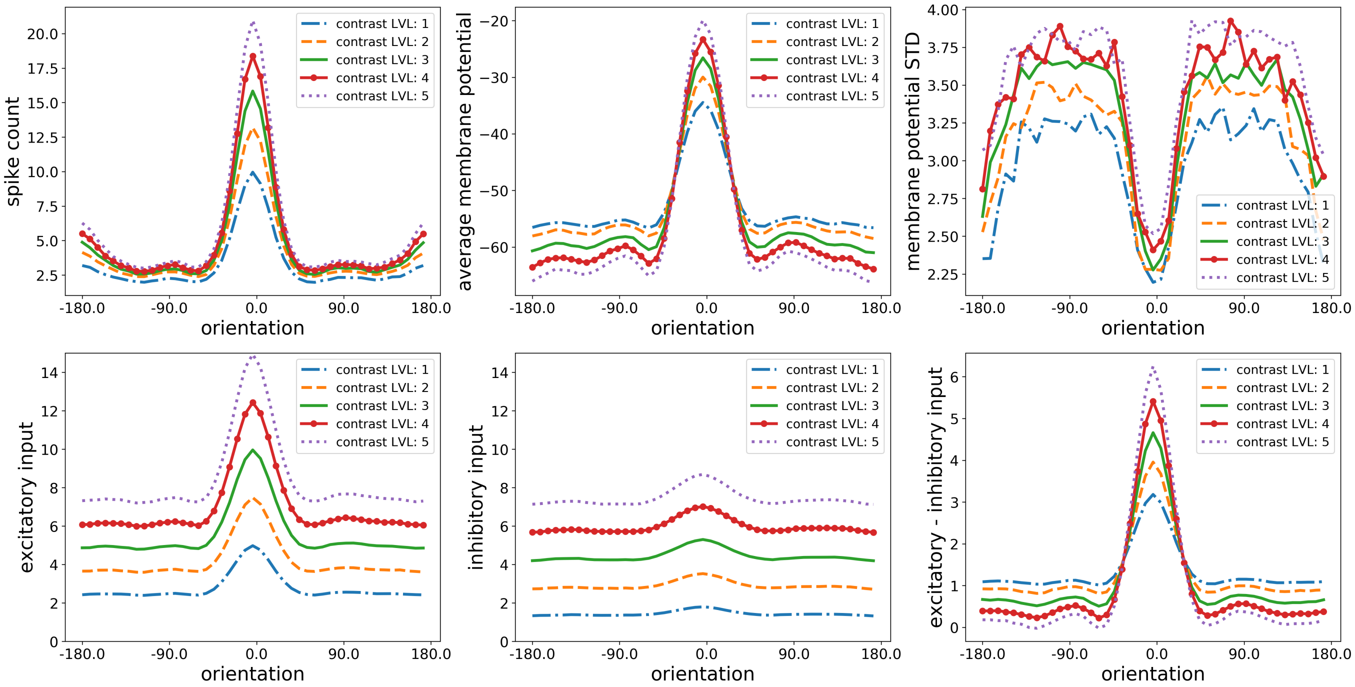


S 7: **Orientation tuning as a function of input contrast, $EI2/1$ model**. Mean spike count (upper left), average membrane potential (upper middle), standard deviation (upper right), mean excitatory input (lower left), mean inhibitory input (lower middle), and difference between excitation and inhibition (lower right).

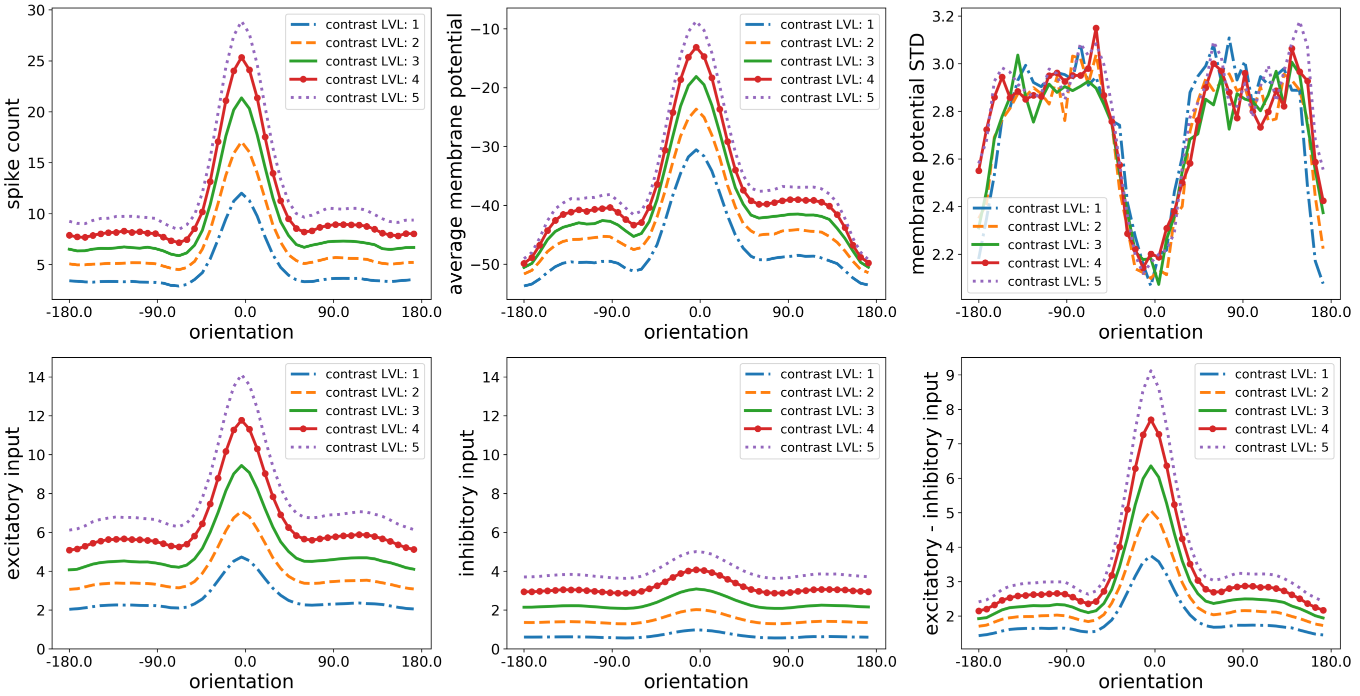


S 8: **Orientation tuning as a function of input contrast, $EI3/1$ model**. Mean spike count (upper left), average membrane potential (upper middle), standard deviation (upper right), mean excitatory input (lower left), mean inhibitory input (lower middle), and difference between excitation and inhibition (lower right).

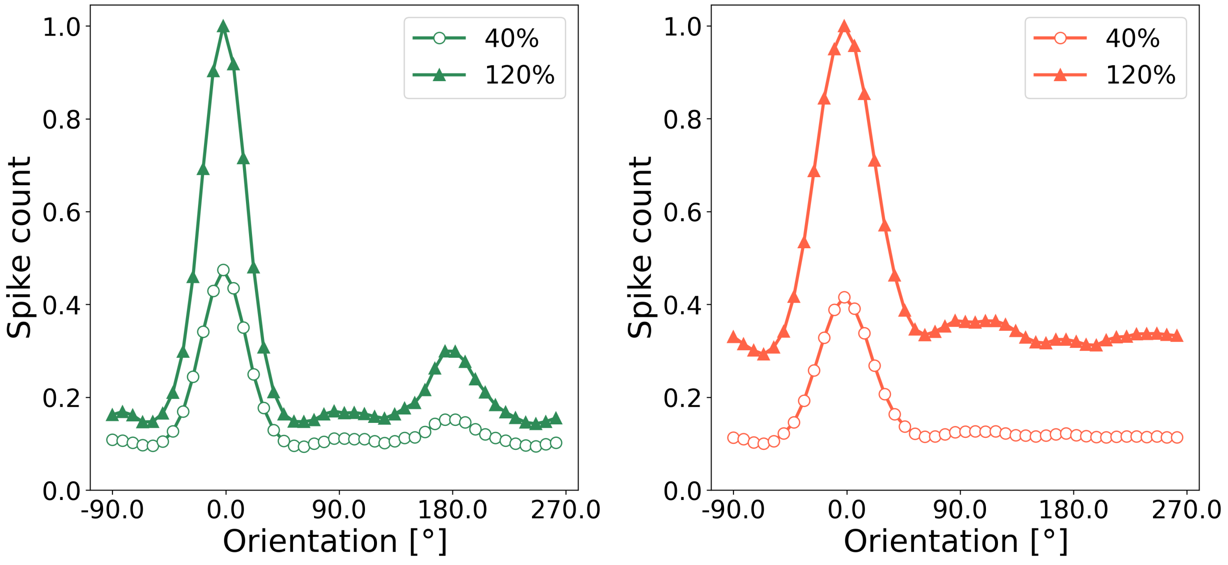


(a) (b)

S 9: **Normalized tuning curves** Tuning curves are normalized with the maximum spike count on high contrast. **(a)** for $EI2/1$ model, **(b)** the $EI3/1$ model, Mean and standard deviation calculated across the excitatory population.

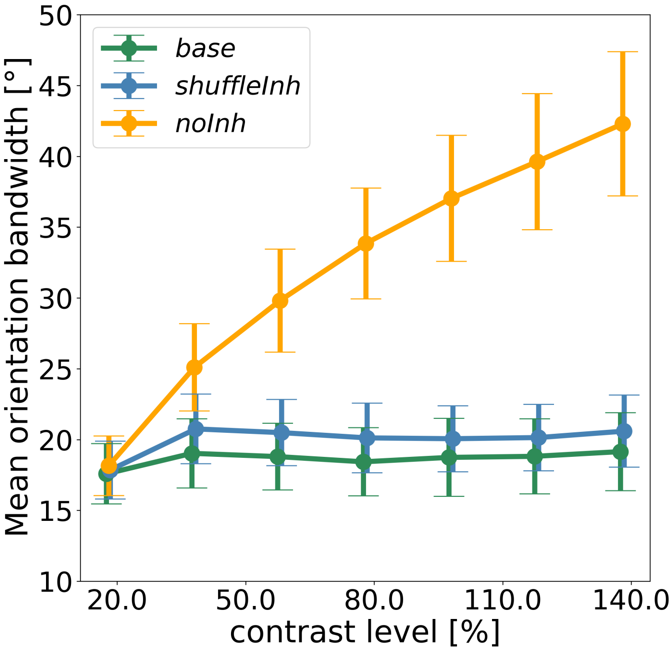


S 10: Mean orientation bandwidth of the excitatory population for different contrast levels. Green: $EI2/1$ model. Orange: Deactivated inhibition, blue: Randomly shuffled feed-forward and feedback inhibition.

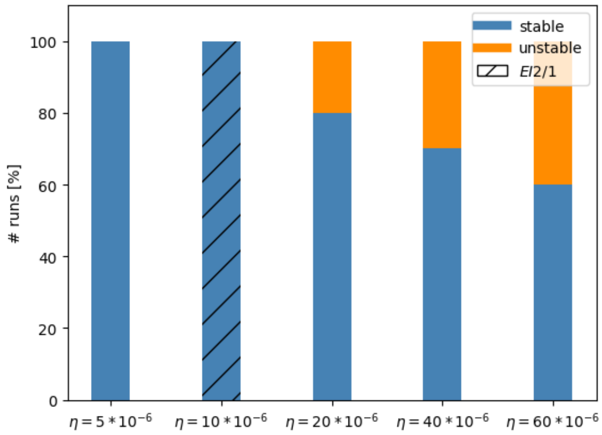


(a)

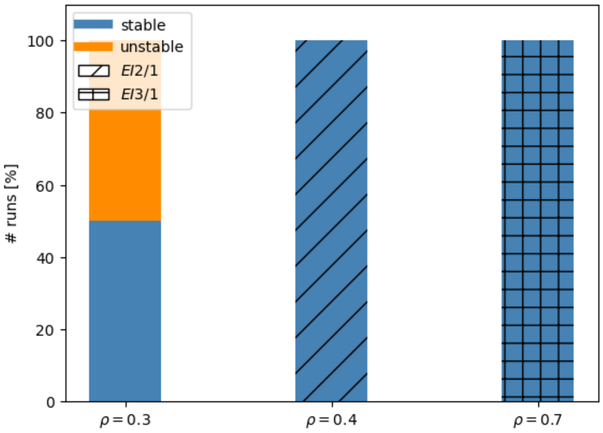


(b)

S 11: **(a)** Percent of runs in which stable receptive fields or unstable (eliminated) receptive fields emerged during learning for different values of $\eta$ (learning rate) of the Vogels et al. (2011) learning rule. Other parameters are taken from the $EI2/2$ model configuration. **(b)** Percent of runs, where stable receptive fields or unstable (eliminated) receptive fields emerged during learning for different $\rho$ (postsynaptic target rate) of the Vogels et al. (2011) learning rule. Other parameters are taken from the $EI2/2$ model configuration. Please note, that $\rho = 0.7$ corresponds to the $EI3/1$ model.

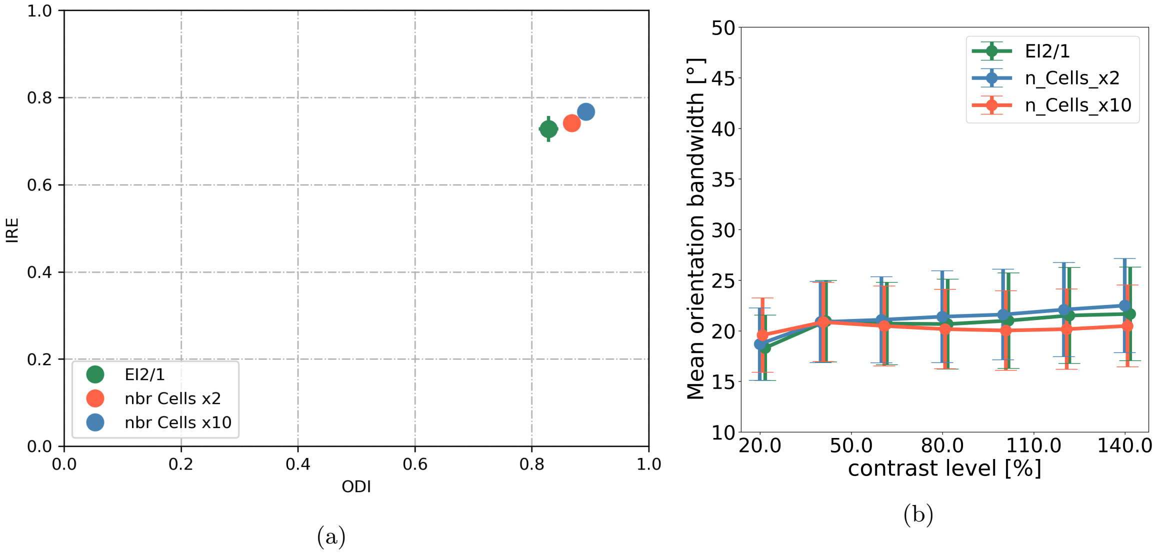


S 12: **Different sized excitatory and inhibitory populations. (a)** Image reconstruction error (IRE) as a function of orientation diversity. **(b)** Orientation Bandwidth (OBW) for different contrast levels. Data from the $EI2/1$ model (green), with twice the number of neurons (red) and with ten times the number of neurons (blue). Note: The number of inhibitory neurons is chosen to fit the $4:1$ excitation to inhibition ratio.

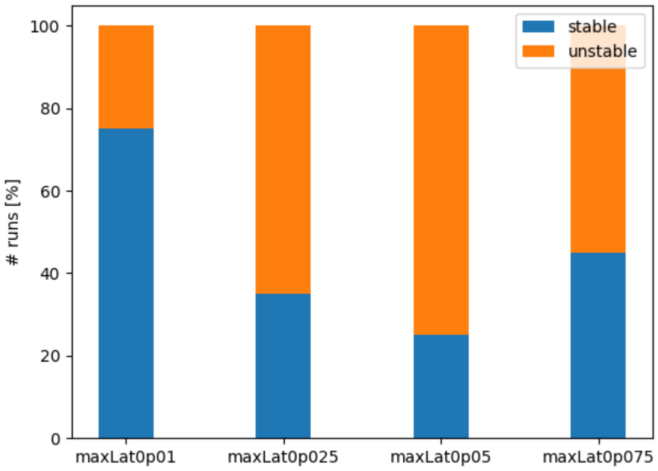


(a)

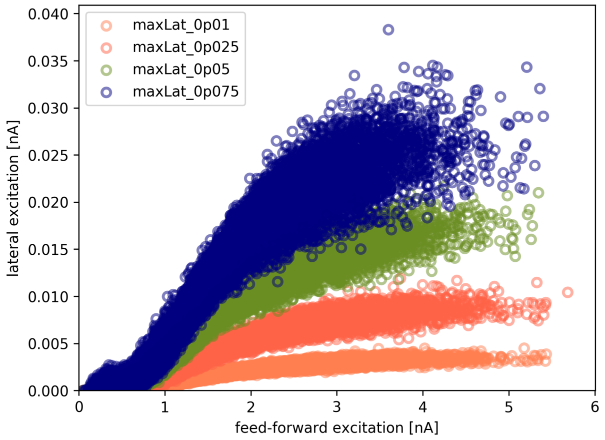


(b)

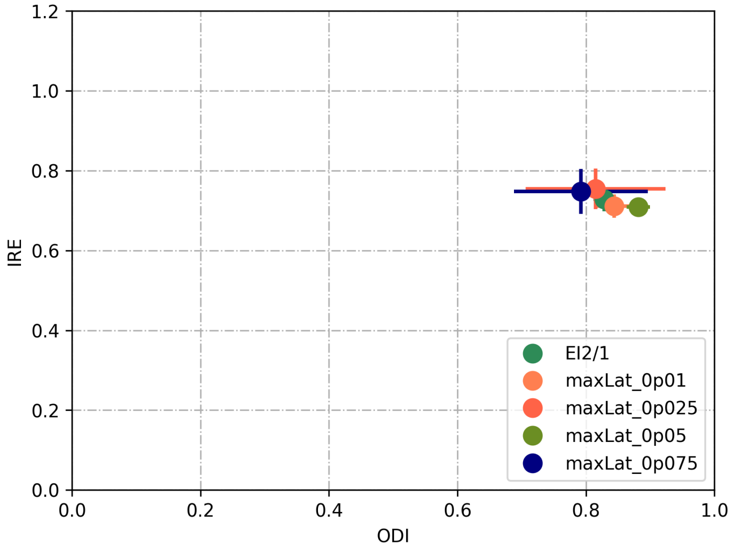


(c)

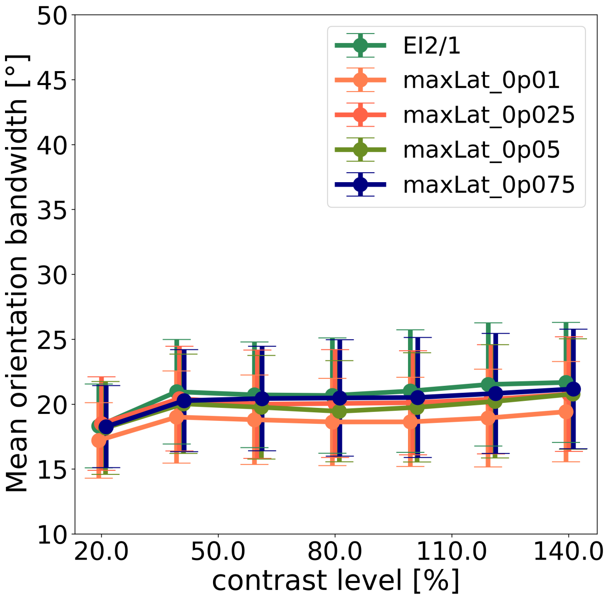


(d)

S 13: **Weak lateral excitation** Recurrent weights are chosen randomly from a normal distribution with $\mu = 0$ and different values of $\sigma$ to control the maximum weight. Negative weight values are set to zero. Blue indicates a maximum weight value of 0.075, olive green indicates a maximum weight value of 0.05, red indicates a maximum weight value of 0.025 and orange indicates a maximum weight values of 0.01. Dark green indicates the $EI2/1$, which is presented for comparison. (a) Percentage of simulations where learning was successful (stable receptive fields emerged) and not successful (all weights in the network run against the maximum weight values). (b) Recurrent excitatory input current as a function of the excitatory current over feed-forward synapses. (c) IRE as a function of the ODIe. (d) OBW for sinusoidal gratings on different levels of contrast.

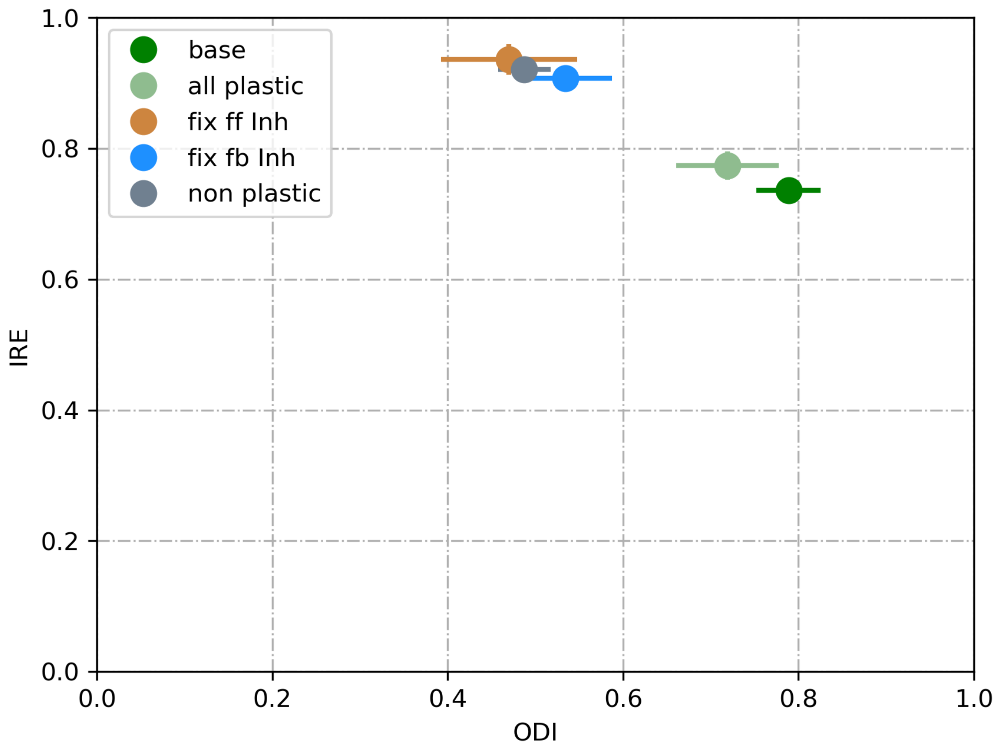


S 14: Image reconstruction error (IRE) as a function of orientation diversity. Excitatory synapses learned with the Pfister & Gerstner (2009) STDP learning rule. Points mark the mean values and the whiskers the standard deviation across 10 model runs.

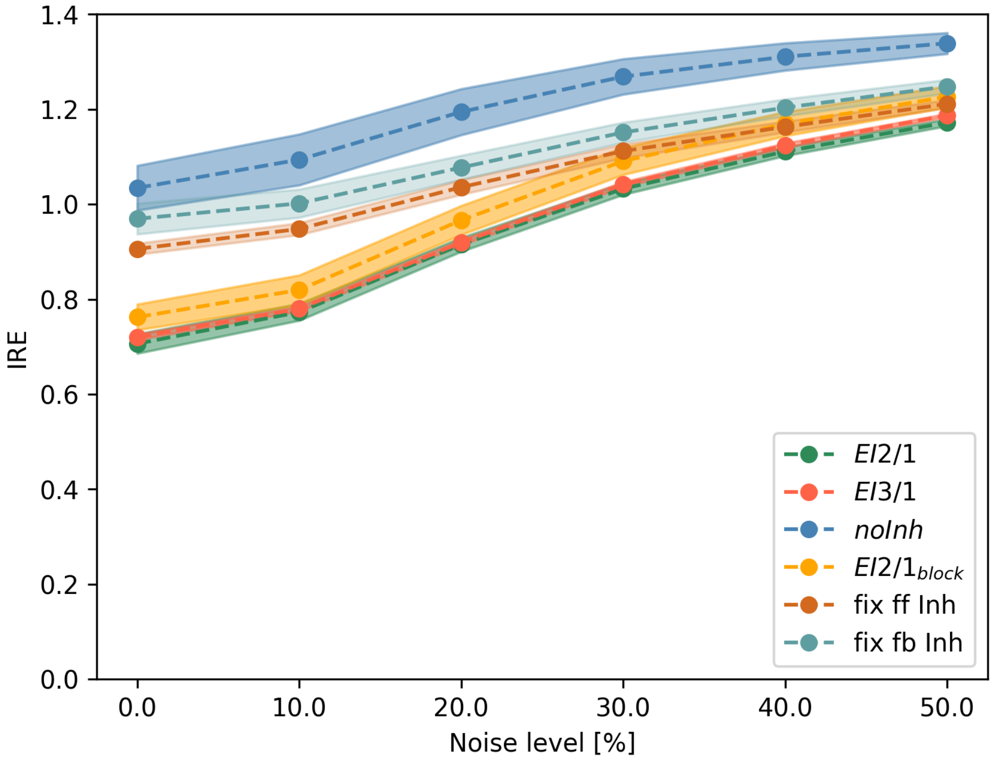


S 15: Image reconstruction error (IRE) as a function of the strength of withe noise. Noise is generated via a normal distribution and added to the natural scene input. The strength is in relation to the maximum pixel value of the original input. Values showing the average IRE of 20 runs for each model configuration, the shaded area represents the standard deviation.
